# *DESpace*: spatially variable gene detection via differential expression testing of spatial clusters

**DOI:** 10.1101/2023.04.17.537189

**Authors:** Peiying Cai, Mark D Robinson, Simone Tiberi

## Abstract

**Motivation:** Spatially resolved transcriptomics (SRT) enables scientists to investigate spatial context of mRNA abundance, including identifying spatially variable genes (SVGs), i.e., genes whose expression varies across the tissue. Although several methods have been proposed for this task, native SVG tools cannot jointly model biological replicates, or identify the key areas of the tissue affected by spatial variability.

**Results:** Here, we introduce *DESpace*, a framework, based on an original application of existing methods, to discover SVGs. In particular, our approach inputs all types of SRT data, summarizes spatial information via spatial clusters, and identifies spatially variable genes by performing differential gene expression testing between clusters. Furthermore, our framework can identify (and test) the main cluster of the tissue affected by spatial variability; this allows scientists to investigate spatial expression changes in specific areas of interest. Additionally, *DESpace* enables joint modelling of multiple samples (i.e., biological replicates); compared to inference based on individual samples, this approach increases statistical power, and targets SVGs with consistent spatial patterns across replicates. Overall, in our benchmarks, *DESpace* displays good true positive rates, controls for false positive and false discovery rates, and is computationally efficient.

**Availability and implementation:** *DESpace* is freely distributed as a Bioconductor R package.

## 1 Introduction

Spatially resolved transcriptomics (SRT) technologies allow the spatial characterization of gene expression profiles in a tissue. SRT techniques can be broadly grouped into two categories: sequencing-based methods (e.g., Slide-seq [28], Slide-seq-V2 [33], 10X Genomics Visium [1], spatial transcriptomics [34], high definition spatial transcriptomics [37], STEREO-seq [5]), and in situ hybridization technologies (e.g., seqFISH [10], seqFISH2 [30], seqFISH+ [9], and MERFISH [14]). Sequencing-based approaches provide measurements across the entire transcriptome but the mRNA abundances (typically) refer to the aggregation of multiple cells; conversely, the imaging-based methods are targeted towards a (usually) limited number of genes but offer higher spatial resolution, e.g., molecular-level measurements. In both cases, computationally efficient methods are required to deal with an increasingly large number of genes and spatial measurements.

The emergence of SRT technologies has prompted the development of novel analysis frameworks that exploit the joint availability of gene expression and spatial information; in particular, below, we briefly introduce two such schemes: spatially resolved clustering and spatially variable gene identification. Other analyses include strategies to identify putative cell-cell communication via local ligand-receptor co-expression, and enrichment of cell type interactions [23].

While single-cell RNA sequencing enables clustering cells based on transcriptional profiles, SRT allows the identification of spatial clusters (Figure 1). Such structures are typically estimated via spatially resolved clustering tools, such as *BayesSpace* [40], *StLearn* [21], *Giotto* [27], and *PRECAST* [15], which cluster spots based on both their gene expression and spatial localization; alternatively, spatial clusters can also be obtained via manual annotation, e.g., from histology [19]. Another popular analysis performed on SRT data consists in identifying genes whose expression patterns change across the tissue, also known as spatially variable genes (SVGs). In particular, one can distinguish between spatial variability (SV) across the entire tissue, and SV within specific areas, such as spatial clusters. The former is arguably the classical SVG application, and is the focus in this manuscript; the latter, instead, targets more subtle variations that could also be of interest. While most SVG methods perform the first task, only a few tools enable studying the second one; for instance, when spatial structures are provided, *nnSVG* [38] can detect both types of SVGs.

**Figure 1:**
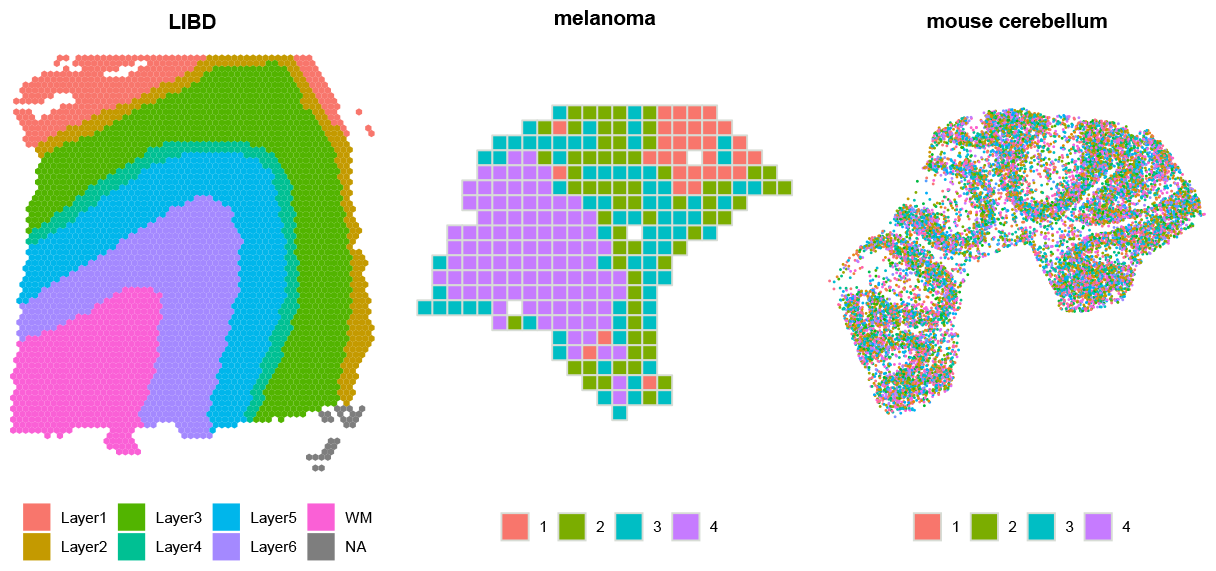
Spatial clusters in three samples from *LIBD* [19] (Visium [1]), *melanoma* [35] (spatial transcriptomics [34]) and *mouse cerebellum* [4] (Slide-seqV2 [33]) datasets. Spatial clusters were obtained: via manual annotations form a pathologist (*LIBD*), *BayesSpace* [40] (*melanoma*), and *StLearn* [21] (*mouse cerebellum*).

In recent years, various methods have been proposed to discover SVGs; notably: *MERINGUE* [3], a density-independent approach built on the spatial auto-correlation of each gene; *nnSVG* [38], which fits nearest-neighbor Gaussian processes; *SpaGCN* [11], that identifies spatial domains based on a graph convolutional network, and performs a Wilcoxon rank-sum test on the mean-level expression changes between domains; *SPARK* [31], using generalized linear mixed models with overdispersed Poisson and Gaussian distributions; *SPARK-X* [32], from the authors of *SPARK*, based on a scalable non-parametric framework to test the dependence between gene expression covariance and distance covariance; *SpatialDE* [36], which infers the dependency between space and gene expression based on a non-parametric Gaussian process; *SpatialDE2* [13], that applies a Bayesian hidden Markov random field to segment spatial regions and a count-based likelihood to model the variance components of each gene within identified regions; and, *Trendsceek* [7], which evaluates significant spatial gene expression heterogeneity via marked point processes.

Despite the abundance of native SVG tools, there are some limitations with the current approaches, including that: i) most methods are computationally demanding, which is particularly troublesome for recent SRT technologies, such as Slide-seq-V2, which provide hundreds of thousands of measurements per gene; ii) biological replicates are not allowed, and only individual samples can be processed, exposing results to a high degree of (biological and technical) variability; iii) few tools (such as *nnSVG*) can incorporate information about spatial structures of interest, such as spatial clusters; and, iv) it is only possible to test the entire tissue for spatial variability, while scientists cannot perform SV testing on specific regions of interest (e.g., white matter in brain cortex).

Additionally, differential gene expression (DGE) methods have also been been applied to discover differentially abundant genes across human brain regions [19]. In particular, Maynard *et al*. [19] propose pseudo-bulk summaries by cluster (i.e., overall abundance across all spots in a cluster) with differential testing between clusters (via *limma* [25]), looking for any change based on the t- or ANOVA F-statistics. Corresponding wrapper functions are provided in the *spatialLIBD* R package [20]. This procedure is computationally efficient, can incorporate multiple samples, and identifies the key clusters affecting SV (via the enrichment test). However, due to its pseudo-bulk nature, it requires biological replicates, and cannot run on individual samples.

Alternatively, one could follow a similar approach, that does not require multiple samples, by using marker gene methods across spatial structures, such as *scran*’s *find-Markers* [17] and *Seurat*’s *FindAllMarkers* [29]. These tools are fast, and can test specific clusters. Nonetheless, they are designed to identify marker genes in each cluster, therefore (for every gene) they perform a statistical test on each individual cluster; these results have to be aggregated if one is interested in studying SVGs across the entire tissue.

Here, we propose *DESpace*, which is a two-step framework based on spatial clustering, and on differential expression testing across clusters. Our approach requires pre-computed spatial clusters (Figure 1), which can be obtained via manual annotation or from spatially resolved clustering tools (e.g., *BayesSpace* [40], *StLearn* [21], Giotto [27], and PRECAST [15]); these spatial clusters are used as a proxy for the spatial information. We then fit a negative-binomial model, which was recently shown to accurately model SRT data [2, 41], via *edgeR* [6,8,18], a popular tool for differential gene expression; we use spatial clusters as covariates, and perform a differential gene expression test across these clusters. If the expression of a gene is significantly associated to the spatial clusters, its expression varies across the tissue, thus indicating a SVG. Albeit our approach resembles marker gene methods and *SpatialLIBD*’s wrappers, it differs from both. Unlike marker methods, that generally identify changes in each individual cluster (i.e., 1 test per cluster), we aim at overall changes across all spatial clusters (i.e., 1 test across all clusters). Furthermore, while *SpatialLIBD*’s approaches compute pseudo-bulk values, our framework models individual (spot-level) measurements; as a consequence, *DESpace* does not require biological replicates, and can also run on individual samples. Clearly, our approach relies on spatial clusters being available, and accurately summarizing the spatial patterns of transcription, but we argue that these assumptions are usually fulfilled. In fact, even when pre-computed annotations are not available, spatially resolved clustering tools enable inferring clusters that summarize the spatial structure of gene expression.

The approach we propose has several advantages. In our benchmarks (see Results), *DESpace* displays good sensitivity and specificity, and is computationally efficient. Furthermore, our two-step framework: i) can jointly model multiple samples, hence increasing statistical power, reducing the uncertainty that characterizes inference performed from individual samples, and identifying genes with coherent spatial patterns across biological replicates; ii) allows identifying the main cluster of the tissue affected by SV, testing if the average expression in a particular region of interest (e.g., cancer tissue) is significantly higher or lower than the average expression in the remaining tissue (e.g., non-cancer tissue), hence enabling scientists to investigate changes in mRNA abundance in specific areas which may be of particular interest. Finally, our tool is flexible, and can input any type of SRT data.

## 2 Materials and methods

### 2.1 Inference with *DESpace*

Consider a SRT dataset, from a single sample, with *N* spots, and where a total of *C* spatial clusters have been identified; further assume that sponds to *N*_*c*_ spots, where 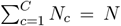. We define *x*_*gi*_ as the mRNA abundance for the *g*-th gene in the *i*-th spot belonging to the *c*-th cluster; we model this value with a negative binomial distribution:

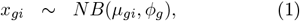

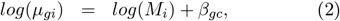

where *NB*(*µ, ϕ*) denotes a negative binomial random variable with mean *µ* and variance *µ*(1 + *µϕ*); *ϕ*_*g*_ is the gene-specific dispersion parameter modeling the variability between spots; *M*_*i*_ is the effective library size (total count multiplied by TMM normalization factor [26]) for the *i*-th spot; and *β*_*gc*_ is the *c*-th spatial cluster coefficient for gene g, which represents the abundance of gene *g* in spatial cluster *c*.

In order to investigate if gene *g* is spatially variable, we verify if its gene expression varies across spatial clusters. In particular, we employ a likelihood ratio test (LRT) [39] to test whether the *β*_*gc*_ parameters differ across clusters, via the following system of hypotheses:

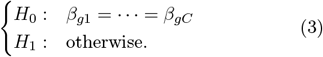

In order to identify the specific regions affected by spatial variability, for each gene, we also apply a LRT on each individual spatial cluster parameter, *β*_*gc*_.

When biological replicates are available, we add a sample-specific parameter to the mean of the negative binomial model in (1). In particular, assuming that the observation comes from the *j*-th replicate, equation (2) becomes:

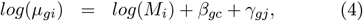

where *γ*_*gj*_ indicates the coefficient for the *j*-th sample in gene *g*, with *γ*_*g*1_ = 0 for the first sample (taken as baseline) to ensure identifiability. SVGs are identified, as in the individual-sample case, by testing for the significance of *β*_*gc*_’s parameters.

In all cases, parameter inference and differential testing are performed via *edgeR* [6, 8, 18]. Note, that we also tried alternative approaches for differential gene expression, namely *DESeq2* [16] and *limma* [25], but overall performance (in terms of sensitivity, specificity, and run-time) was better when using *edgeR*.

### 2.2 Simulation studies - anchor data

We have designed several benchmarks, on real and semi-simulated data; all our analyses start from one of three SRT datasets (collected from distinct spatial technologies), denoted by *LIBD* [19], *melanoma* [35] and *mouse cerebellum* [4] (Figure 1).

The *LIBD* [19] data, collected via the Visium platform [1], contains 12 dorso-lateral pre-frontal cortex samples from 3 independent human brain donors (i.e., biological replicates). In all samples, the tissue was partitioned via manual annotation from pathologists into white matter and up to six layers (Figure 1). Overall, this dataset contains, for each sample, on average, measurements for 33,538 genes across 3,973 spots; after filtering (see Supplementary Details), this reduces to 14,628 genes and 3,936 spots (Supplementary Table 1).

The *melanoma* [35] data consists of 8 melanoma lymph node samples from 4 distinct individuals (i.e., biological replicates), diagnosed with stage IIIc melanoma. This data was obtained by applying spatial transcriptomics (ST) [34] technology to melanoma lymph node biopsies. In the original study [35], the spots of each sample were clustered in four groups: melanoma, stroma, lymphoid tissue, and remaining spots. We clustered spots in four spatial clusters using *BayesSpace* (Figure 1), where cluster 4 indicates the melanoma area, cluster 2 embeds both stroma and lymphoid tissues, cluster 3 represents the border region between lymphoid and tumor tissues, and cluster 1 includes lymphoid tissue distant from the cancer. The dataset, on average, consists of a total of 15,884 genes per sample, measured on 294 spots; after filtering (see Supplementary Details), we retained 8,173 genes and 290 spots (Supplementary Table 1).

The *mouse cerebellum* [4] data, obtained via Slide-seqV2 [33] technology, consists of measurements, from a single sample, for 20,141 genes across 11,626 beads. After filtering (see Supplementary Details), these numbers reduced to 8,277 and 11,615, respectively (Supplementary Table 1). Beads were clustered via *StLearn* (Figure 1), obtaining similar clusters to those computed by Zhu *et al*. (2021) [32], on the same dataset, using RCTD software [4].

Note that for the semi-simulated data from the *LIBD* and *melanoma* datasets, we generated one simulation for each biological replicate (3 and 4 in the former and latter case, respectively).

### 2.3 Simulation studies - SV profiles

We generated various semi-simulated datasets, by using, as anchor data, the previously described three datasets. In particular, we edited the *runPatternSimulation* function from *Giotto* [27], and re-arranged the real measurements to partition the space into a highly and a lowly abundant region (see Supplementary Details); by partitioning the tissue in different ways, we generated five different SV patterns. We initially separated the space in artificial regions: top *vs*. bottom (or right *vs*. left) areas (Figure 2, *Bottom/Right*), and inside *vs*. outside a circle (Figure 2, *Circular*). Then, we followed real data structures such as brain cortex layers and melanoma regions (Figure 2, *Annotations*). In these three scenarios, in each simulated dataset, half of the SVGs are highly abundant in one region (e.g., bottom or centre of the circle), and half are highly abundant in the complementary region (e.g., top or outside the circle). However, in the cases described above, all SVGs follow the same spatial structures, which is unlikely to happen in real data. In order to improve the realism of the semi-simulated data, we designed two additional simulations, where SVGs follow multiple spatial patterns, all based on real data structures; we generated two such structures, referred to throughout as *mixture* and *inverted mixture*. In the former case, a small region is identified as highly abundant, while most of the tissue is lowly abundant; conversely, in the latter case, gene expression is uniform in most of the tissue, and a small region is characterized by low abundance (Supplementary Figure 1). In each simulation, we generated between 33% (*bottom/right, circular*, and *annotations* patterns) and 50% (*mixture*, and *inverted mixture* patterns) of SVGs, while the remaining genes are uniformly distributed; such uniform patterns were obtained by randomly permuting measurements across the tissue (see Supplementary Details).

**Figure 2:**
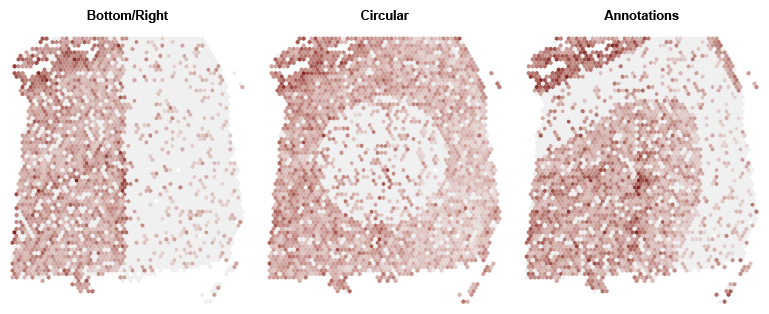
Three examples of simulated SVGs, from the *LIBD* data, following *bottom/right, circular*, and *annotations* patterns. Examples of SVGs from *mixture* and *inverted mixture* patterns are presented in Supplementary Figure 1.

Note that, when running *DESpace* and marker methods, we did not use the original spatial clusters that we simulated from. Instead, we employed noisy spatial clusters inferred via *StLearn* and *BayesSpace*. We choose these two spatially resolved clustering tools, because they performed well in recent benchmarks [40].

## 3 Results

### 3.1 Simulation studies - individual sample

In all simulations, we bechmarked *DESpace* against seven of the most popular tools for SVG detection: *SpaGCN* [11], *SpatialDE* [36], *SpatialDE2* [13], *SPARK* [31], *SPARK-X* [32], *MERINGUE* [3], and *nnSVG* [38]. Additionally, we considered two tools originally designed to detect marker genes: *scran’s findMarkers* [17] and *Seurat*’s *FindAllMarkers* [29]; similarly to *DESpace*, both methods were applied to spatial clusters computed by *StLearn* and *BayesSpace*. Note that the *limma* wrappers in *SpatiaLIBD* were not considered here, because the approach requires multiple samples.

Figure 3 reports the true positive rate (TPR) *vs*. false discovery rate (FDR) of all methods for the three datasets and five spatial patterns. Rates for the *LIBD* and *melanoma* samples are aggregated across biological replicates; the results for the individual samples can be seen in Supplementary Figures 2-3). In all five SV patterns *DESpace* controls for the false discovery rate, and, in most cases, leads to a higher statistical power than competitors; interestingly, this gap is smaller (or absent) in simpler SV patterns, and increases in the more complex ones. This is particularly true for *scran*’s *find-Markers*, which behaves similarly to *DESpace* in the *bottom/right, circular* and *annotations* simulations, all defined by 2 clusters only, while displays a loss of power in the *mixture* and *inverted mixture* cases, that are characterized by 3 or more clusters. This is possibly due to the fact that marker methods detect differences between pairs of clusters, and, are not designed to identify global changes across three or more clusters. Overall, these findings suggest that our framework may be particularly beneficial for detecting challenging spatial structures. Notably, results are highly consistent across the three datasets, and *DESpace* leads to similar rates when inputting spatial clusters estimated from *StLearn* or *BayesSpace*, which indicates that our approach is robust with respect to the spatial clusters provided.

**Figure 3:**
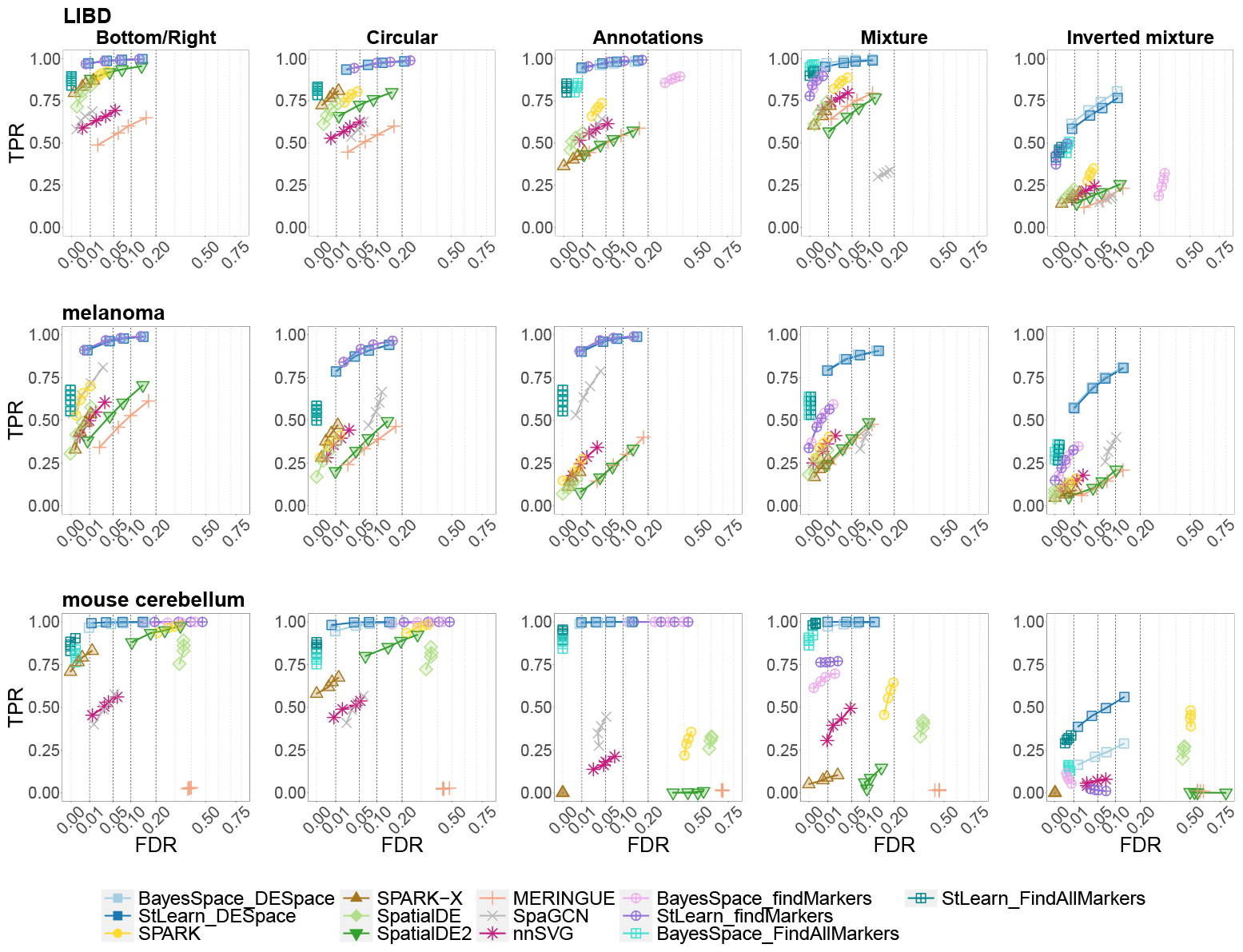
TPR vs. FDR for SVG detections in the individual sample simulations. Rows and columns refer to the anchor data used in the simulation, and to the SV profiles, respectively. *BayesSpace_DESpace, BayesSpace_findMarkers*, and *BayesSpace_FindAllMarkers*, as well as their counterparts *StLearn_DESpace, StLearn_findMarkers*, and *StLearn_FindAllMarkers*, indicate *DESpace, scran*’s *findMarkers*, and *Seurat*’s *FindAllMarkers*, respectively, based on spatial clusters computed via *BayesSpace* and *StLearn*.

Furthermore, in order to investigate false positive detections, in each simulation, we selected the subgroup of uniform genes (i.e., non-SVGs), and studied the distribution of raw p-values. Overall, most methods (including *DESpace*) display approximately uniform p-values, and none presents a significant inflation of false positives, with the only exception of *SPARK* on the *mouse cerebellum* data, and *FindAllMarkers* on all datasets (Supplementary Figure 4). Notably, *SpatialDE* and *nnSVG* produce highly conservative results, with p-values significantly inflated towards 1, which is consistent with previous findings [31, 38].

Additionally, we investigated if methods are able to sort SVGs according to the strength of their spatial structure. In fact, it is desirable that genes displaying strong spatial patterns are ranked before (i.e., smaller p-value) genes with weaker spatial patterns, because the first ones are more likely to be biologically relevant. To this aim, we generated a simulation with SVGs only, where 50% of the genes display a strong spatial pattern, and 50% display a weak one (see Supplementary Details). Supplementary Figure 5 displays the TPRs for the strong and weak patterns, while Supplementary Tables 4-5 report the respective areas under the curves; overall, *DESpace* and marker methods rank strong SV patterns better than competitors.

We designed an additional simulation to systematically assess how the number of clusters affects our analysis framework. Specifically, we considered sample 151507 from the *LIBD* data, and artificially partitioned its tissue in 2, 4, 6, 8, 10, and 12 clusters (Supplementary Figure 6); in each case, we simulated a *mixture* pattern with 50% uniform genes and 50% spatial patterns, equally distributed among the clusters. For a homogeneous comparison, we chose exactly the same SVGs acoss scenarios. We then fit *DESpace* using the clusters estimated via *BayesSpace* and *StLearn* (shown in Supplementary Figures 7-8); additionally, we also fit *DESpace* on the original clusters the data was simulated from (as in Supplementary Figure 6). In all scenarios, the FDR is well calibrated. Furthermore, when using the original spatial clusters, the TPR is also stable (Supplementary Figure 9, left panel). Conversely, when using noisy estimated clusters from *BayesSpace* and *StLearn*, the statistical power marginally decreases as the number of clusters increases (particularly when using *StLearn*) (Supplementary Figure 9, middle and right panels). Therefore, while increasing the number of clusters does not appear to impact *DESpace* directly, it leads to more noisy clusters estimated by spatial clustering tools, which in turn may affect the TPR of our framework.

### 3.2 Simulation studies - individual cluster

We also investigated the ability of our two-step framework to target the main cluster affected by SV. Note that, when only 2 spatial clusters are available, individual cluster testing corresponds to the global test described above; therefore, this analysis framework can only be applied when at least 3 clusters are available, and is particularly beneficial when there is a large number of clusters. To validate this approach, we considered the *mixture* and *inverted mixture* simulations in the *LIBD* dataset, which contain between 5 and 7 spatial clusters per biological replicate. In these simulations, it is possible to define a key spatial cluster that displays the biggest difference compared to the average signal; in the *mixture* (or *inverted mixture*) simulations this corresponds to the small region with higher (or lower) abundance than the majority of the tissue (Supplementary Figure 1). Here, we compared our framework to marker gene methods only, because native SVGs approaches cannot perform this kind of testing.

On average, when using *BayesSpace* clusters, *DESpace* identified the main SV cluster in 99.6 and 88.9% of SVGs in *mixture* and *inverted mixture* patterns, respectively; using *StLearn* clusters led to similar numbers Table 1). Marker methods performed similarly in the *mixture* simulation, but displayed a lower accuracy in the *inverted mixture* pattern (Table 1, and Supplementary Table 2). Note that, in general, percentages are higher in the *mixture* simulations, because the change in abundance, between the key spatial cluster and the rest of the tissue, is higher compared to the *inverted mixture* patterns (Supplementary Figure 1). Furthermore, in non-SVGs, *DESpace* p-values were uniformly distributed in all spatial clusters, showing no inflation of false positive detections at the cluster level, while *findMarkers* displayed conservative results, and *FindAllMarkers*’s p-values, although generally uniform, were occasionally inflated towards 0 (Supplementary Figures 10-13). This testing feature may be particularly useful for computational biologists, as it enables them to directly target specific regions of the tissue, for example by identifying SVGs characterized by high or low abundance in white matter or layer 3 (Figure 4). Finally, note that we also provide a faster implementation of the individual cluster testing, where we recycle the previously-computed dispersion estimates from the gene-level test (see Supplementary Details); this strategy performs similarly to the original one (analogous fraction of key spatial clusters identified), while the average runtime decreases significantly: from 32 to 2 minutes per sample (Supplementary Table 3).

**Table 1:** Individual cluster results. Percentage of times that *DESpace, Seurat*’s *FindAllMarkers* and *scran*’s *findMarkers* identified the main SV cluster in *mixture* (i.e., Mixture) and *inverted mixture* simulations (i.e., Inverted), using *BayesSpace* and *StLearn* clusters. Percentages are averages across the three samples; sample specific results are visible in Supplementary Table 2.

| Pattern | *DESpace* |  | *FindAllMarkers* |  | *findMarkers* |  |
| --- | --- | --- | --- | --- | --- | --- |
|  | *BayesSpace* | *StLearn* | *BayesSpace* | *StLearn* | *BayesSpace* | *StLearn* |
| Mixture | 99.6 | 99.1 | 99.6 | 99.6 | 98.5 | 97.8 |
| Inverted | 88.9 | 84.7 | 84.2 | 82.5 | 60.7 | 79.8 |

**Figure 4:**
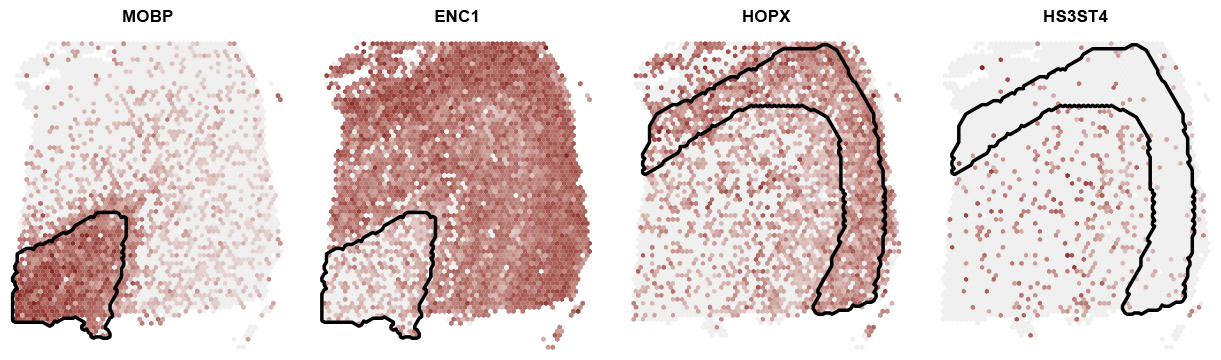
Expression plot, for four SVGs detected with *DESpace* individual cluster test, on the real *LIBD* dataset (sample 151673). SVGs were identifying by selecting high and low expression in white matter (genes MOBP and ENC1, respectively), and high and low abundance in layer 3 (genes HOPX and HS3ST4, respectively). Lines highlight the cluster being tested.

### 3.3 Simulation studies - multiple samples

Using the *LIBD* and *melanoma* datasets, which contain three and four biological replicates, respectively, we designed a multi-sample simulation: we simulated the 5 SV profiles shown before, and ensured that SVGs (and their spatial profiles) are consistent across samples; we then fit *DESpace* on, both, single-sample and multi-sample modes. Here, we aim to compare the two *DESpace* approaches, based on identical input data; therefore, in this analysis, we used the same spatial clusters (i.e., the annotations we simulated from) for both single-sample and multi-sample modes. While both approaches control for the FDR, jointly modelling multiple samples leads, in all cases, to a significant increase of the TPR, which is expectable given that more information is available (Supplementary Figures 14-16).

Additionally, we compared our multi-sample approach against alternative tools that also allow the joint modelling of multiple samples; in particular, we included *SpatialLIBD*’s *limma* wrappers, as well as marker gene methods. Note that *SpatialLIBD*’s ANOVA approach was only applied to the *mixture* and *inverted mixture* simulations, because it requires 3 or more clusters. Native SVG methods were excluded because they do not allow joint modelling of multiple samples. Compared to *SpatialLIBD*’s wrappers and marker methods, our multi-sample approach behaves similarly when there are two clusters only, while it displays a higher TPR in the presence of three or more clusters (i.e., in *mixture* and *inverted mixture* simulations) (Figure 5).

**Figure 5:**
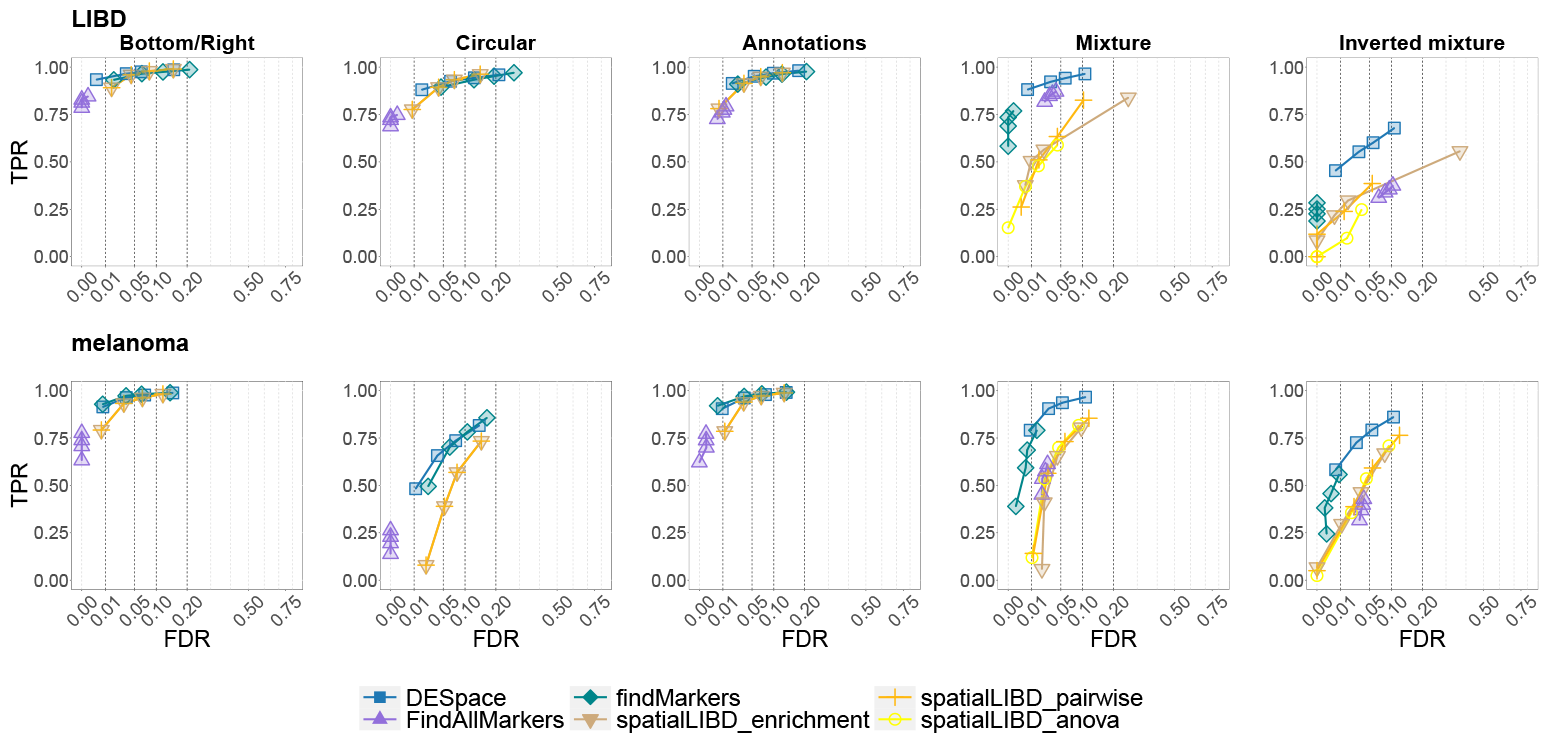
TPR vs. FDR for SVG detections in the multiple sample simulation. Rows and columns refer to the anchor data used in the simulation, and to the SV profiles, respectively. Note that TPRs are lower than in the individual simulation (Figure 3), because we have simulated slightly weaker spatial patterns here (see Supplementary Details).

Furthermore, note that, when sample-specific information is available (e.g., batch effects), it can be incorporated in *DESpace* in the form of additional covariates for the mean abundance in (2). DGE testing is then performed on spatial clusters only, to identify SVGs.

### 3.4 Applications to real data

We also applied all methods to the three real datasets considered. For the *LIBD* and *melanoma* datasets, we considered all 12 and 8 samples, respectively. In all datasets, we computed spatial clusters via *BayesSpace* and *StLearn*; in the *LIBD* dataset, we additionally considered pre-computed manually annotated clusters. In all cases, *DESpace* identifies significantly more SVGs (at 1% FDR threshold) than any other method, which is coherent with the higher statistical power displayed in simulations.

In *LIBD* and *melanoma* data, we also investigated the coherency of the top ranked genes across different replicates. In order to do that, we used the Jaccard index [12], which ranges between 0 and 1, and measures the similarity between two pairs of groups. In particular, for every method, we considered the top 1,000 (LIBD) or 200 (melanoma) ranked genes (i.e., smallest p-value) in each sample, and computed the Jaccard index on pairs of samples; we then averaged results across samples. In both datasets, *DESpace, FindAllMarkers, SPARK* and textitSPARK-X display the highest Jaccard index, which indicates a greater coherency of top ranked genes across samples, while *SpaGCN* and *findMarkers* are associated to the lowest values of the index (Figure 6). Note that, the Jaccard index of all methods is significantly lower in the *melanoma* dataset, compared to the *LIBD* data, which is reasonable given the high degree of gene expression variability in cancer.

**Figure 6:**
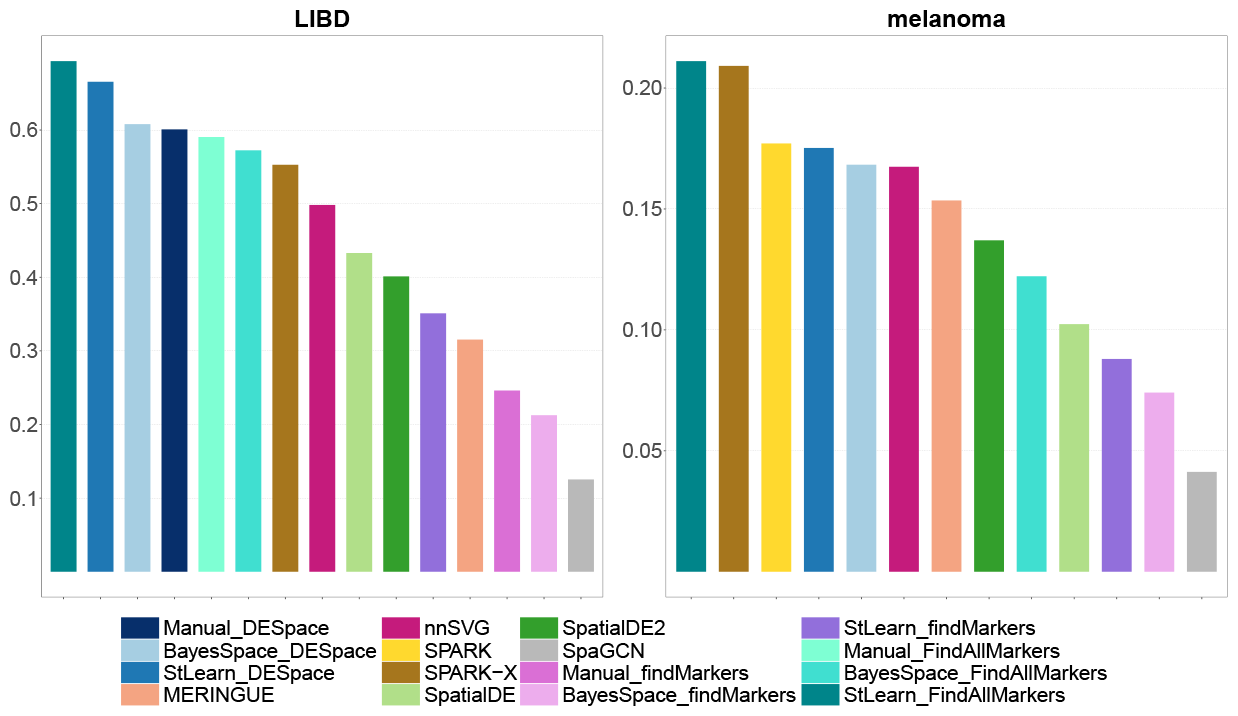
Jaccard index on the *LIBD* and *melanoma* datasets. For each method, the index is measured on the sets of top SVGs reported across replicates, and represents how coherent results are between samples: higher values indicate greater coherency.

In the *melanoma* dataset, we additionally investigated how coherent results are with respect with previous findings. In particular, in absence of a ground truth, we generated three lists of potentially interesting genes based on searches on The Human Protein Atlas website [22]: 19 HLA genes, which are essential for the immune-mediated regression of metastases [35], 12 melanoma marker genes and 3,398 genes associated to the term “melanoma”. We then compared those lists with the top 500 results from every method in each *melanoma* sample. Overall, we found that *DESpace* top discoveries contained more genes from those lists compared to alternative approaches, closely followed by *Seurat*’s *FindAllMarkers* and *SPARK-X* (Supplementary Table 6).

Finally, we investigated the computational efficiency of methods, and computed each tool’s runtime on the three real datasets (Supplementary Figure 17). The ranking of methods is similar across datasets: *SPARK-X, SpaGCN*, and *findMarkers* excel as the fastest methods, followed by *DESpace, FindAllMarkers, SpatialDE* and *SpatialDE2*, while the slowest three methods appear to be *nnSVG, SpaGCN*, and *SPARK*. Note that *DESpace*’s and marker methods’ runtimes largely depend on the computational cost required to generate spatial clusters; in particular, in all datasets, *StLearn* is significantly faster than *BayesSpace*.

## 4 Discussion

In this manuscript, we have presented *DESpace*, an approach based on differential testing across spatial structures to identify spatially variable genes from SRT data. We have run extensive benchmarks on both semi-simulated and real datasets, where we compared our tool to state-of-the-art SVG, marker detection and pseudo-bulk approaches. To the best of our knowledge, this is the first comprehensive benchmark of marker methods for the purpose of SVG detection. Overall, our two-step framework displays good statistical power (i.e., TPR), controls for false positive rates, and is computationally efficient; furthermore, it can i) identify the key area of the tissue affected by SV, and ii) jointly model multiple samples. The former feature allows performing statistical testing on specific regions of the tissue, while the latter one provides increased TPR, and enables the identifications of SVGs with consistent spatial patterns across replicates. In some cases, when only two spatial clusters are present, marker methods perform similarly to *DESpace*; however, our framework has a higher statistical power when three or more clusters are present (i.e., *mixture* and *inverted mixture* simulations), which is the most common scenario in real data.

Additionally, our approach is flexible and can work with SRT data from any technology. Finally, *DESpace* is distributed, open-access, as a Bioconductor R package, which makes it easy to install and integrate with existing pipelines (such as the popular *SpatialExperiment* class [24]), is accompanied by an example vignettes that illustrate its usage, and includes plotting functions that facilitate visualization and interpretation of results.

We also acknowledge some limitations of our approach. In particular, our framework requires pre-computed spatial clusters, and cannot be applied when these are absent; nonetheless, we do not think that this is a barrier in practice, because such structures can be successfully obtained in most datasets via spatially resolved clustering tools. Additionally, *DESpace* is (by design) prone to detecting SVGs that are associated with spatial clusters, and will struggle to identify rare spatial patterns that do not follow these structures. When interest lies in identifying such SVGs, using a native SVG tool will be more appropriate. Furthermore, we are aware of the circularity (or double dipping) issue of our framework; indeed, when relying on spatially resolved clustering tools, we use our data twice: i) to cluster spots, and ii) to perform differential analyses between such clusters. Nonetheless, each individual gene contributes marginally to spatial clustering results, which are (usually) based on the expression data from thousands of genes. In addition, in all our benchmarks, we have empirically shown that circularity does not impact performance results (i.e., good statistical power, and well calibrated false positive rates). Lastly, similarly to the majority of SVG tools, the main application of *DESpace* consists in identifying global spatial structures across the entire tissue. For studying spatial variability within clusters, one could iterate our framework (including spatial clustering) within a specific region of interest, or, alternatively, use a tool specifically designed for this purpose (e.g., *nnSVG*, when providing spatial clusters as covariates).

Finally, a consideration about the computational aspect: recent SRT technologies allow measurements at sub-cellular resolution; this leads to a significant increase in the size of datasets and, in turn, of the computational cost of downstream analyses. In order to mitigate the runtime of our tool, one may aggregate the signal from several spots that belong to the same spatial cluster, which would greatly diminish the number of observations available. We aim to explore this in the future to quantify the reduction in the computational burden, and assess how it affects performance.

## Availability

*DESpace* is freely available as a Bioconductor R package at: https://bioconductor.org/packages/DESpace. The scripts used to run all analyses are available on GitHub (https://github.com/peicai/DESpace_manuscript) and Zenodo (DOI: 10.5281/zenodo.8424676). The *LIBD* [19] data can be accessed via *spatialLIBD* [20] ExperimentHub R Bioconductor packages; the *melanoma* [35] data set is available at https://www.spatialresearch.org/resources-published-datasets/; and the *mouse cerebellum* [4] data can be reached through Broad Institute’s single-cell repository (https://singlecell.broadinstitute.org/single_cell/study/SCP948).

## Acknowledgements

This work was supported by Swiss National Science Foundation grants 310030_175841 and 310030_204869 to MDR. MDR acknowledges support from the University Research Priority Program Evolution in Action at the University of Zurich.

## Author contributions

PC implemented the method, performed all analyses, and contributed to writing the manuscript. MDR contributed to designing the study and to writing the manuscript. ST conceived the method, designed the study and wrote the manuscript. All authors approve the article.

## References

[1] 10x Genomics Acquires Spatial Transcriptomics. https://www.10xgenomics. com/news/10x-genomics-acquires-spatial-transcriptomics., 2018. [Accessed November 2019].

[2] N. BinTayyash, S. Georgaka, S. John, S. Ahmed, A. Boukouvalas, J. Hensman, and M. Rattray. Non-parametric modelling of temporal and spatial counts data from RNA-seq experiments. Bioinformatics, 37(21):3788–3795, 2021.

[3] C. D. X. Z. Brendan F Miller, Dhananjay Bambah-Mukku and J. Fan. Characterizing spatial gene expression heterogeneity in spatially resolved single-cell transcriptomic data with nonuniform cellular densities. Genome research, May 2021.

[4] D. M. Cable, E. Murray, L. S. Zou, A. Goeva, E. Z. Macosko, F. Chen, and R. A. Irizarry. Robust decomposition of cell type mixtures in spatial transcriptomics. Nature biotechnology, 40:517–526, 2022.

[5] A. Chen, S. Liao, M. Cheng, K. Ma, L. Wu, Y. Lai, X. Qiu, J. Yang, J. Xu, S. Hao, X. Wang, H. Lu, X. Chen, X. Liu, X. Huang, Z. Li, Y. Hong, Y. Jiang, J. Peng, S. iu, and J. Wang. High-definition spatial transcriptomics for in situ tissue profiling. Cell, 185(10):1777–1792.e21, 2022.

[6] Y. Chen, A. T. Lun, and G. K. Smyth. From reads to genes to pathways: differential expression analysis of RNA-Seq experiments using Rsubread and the edgeR quasi-likelihood pipeline. F1000Research, 5, 2016.

[7] P. J. Daniel Edsgård and R. Sandberg. Identification of spatial expression trends in single-cell gene expression data. Nature Methods, 15:339–342, 2018.

[8] G. K. S. Davis J McCarthy, Yunshun Chen. Differential expression analysis of multifactor RNA-Seq experiments with respect to biological variation. Nucleic acids research, 40(10):4288–97, 2012.

[9] C.-H. L. Eng, M. Lawson, Q. Zhu, R. Dries, N. Koulena, Y. Takei, J. Yun, C. Cronin, C. Karp, G.-C. Yuan, et al. Transcriptome-scale super-resolved imaging in tissues by RNA seqFISH+. Nature, 568(7751):235–239, 2019.

[10] T. Z. M. A. Eric Lubeck, Ahmet F Coskun and L. Cai. Single-cell in situ RNA profiling by sequential hybridization. Nature methods, 11(4):360–1, 2014.

[11] J. Hu, X. Li, K. Coleman, A. Schroeder, N. Ma, D. J. Irwin, E. B. Lee, R. T. Shinohara, and M. Li. SpaGCN: Integrating gene expression, spatial location and histology to identify spatial domains and spatially variable genes by graph convolutional network. Nature Methods, 18:1342–1351, Novemble 2021.

[12] P. Jaccard. Distribution de la flore alpine dans le bassin des dranses et dans quelques régions voisines. Bull Soc Vaudoise Sci Nat, 37:241–272, 1901.

[13] I. Kats, R. Vento-Tormo, and O. Stegle. SpatialDE2: Fast and localized variance component analysis of spatial transcriptomics. bioRxiv, 2021.

[14] J. R. M. S. W. X. Z. Kok Hao Chen, Alistair N Boettiger. RNA imaging. Spatially resolved, highly multiplexed RNA profiling in single cells. Science (New York, N.Y.), 348(6233), 2015.

[15] W. Liu, X. Liao, Z. Luo, Y. Yang, M. C. Lau, Y. Jiao, X. Shi, W. Zhai, H. Ji, J. Yeong, et al. Probabilistic embedding, clustering, and alignment for integrating spatial transcriptomics data with PRECAST. Nature Communications, 14(1):296, 2023.

[16] M. I. Love, W. Huber, and S. Anders. Moderated estimation of fold change and dispersion for RNA-seq data with DESeq2. Genome biology, 15(12):1–21, 2014.

[17] A. T. Lun, D. J. McCarthy, and J. C. Marioni. A step-by-step workflow for lowlevel analysis of single-cell rna-seq data with bioconductor. F1000Research, 5, 2016.

[18] G. K. S. Mark D Robinson, Davis J McCarthy. edgeR: a Bioconductor package for differential expression analysis of digital gene expression data. Bioinformatics, 26(1):139–140, 11 2009.

[19] K. R. Maynard, L. Collado-Torres, L. M. Weber, C. Uytingco, B. K. Barry, S. R. Williams, J. L. Catallini, M. N. Tran, Z. Besich, M. Tippani, et al. Transcriptome-scale spatial gene expression in the human dorsolateral prefrontal cortex. Nature neuroscience, 24(3):425–436, 2021.

[20] B. Pardo, A. Spangler, L. M. Weber, S. C. Page, S. C. Hicks, A. E. Jaffe, K. Martinowich, K. R. Maynard, and L. Collado-Torres. spatialLIBD: an R/Bioconductor package to visualize spatially-resolved transcriptomics data. BMC genomics, 23(1):434, 2022.

[21] D. Pham, X. Tan, J. Xu, L. F. Grice, P. Y. Lam, A. Raghubar, J. Vukovic, M. J. Ruitenberg, and Q. Nguyen. stLearn: integrating spatial location, tissue morphology and gene expression to find cell types, cell-cell interactions and spatial trajectories within undissociated tissues. BioRxiv, pages 2020–05, 2020.

[22] F. Pontén, K. Jirström, and M. Uhlen. The Human Protein Atlas—a tool for pathology. The Journal of Pathology: A Journal of the Pathological Society of Great Britain and Ireland, 216(4):387–393, 2008.

[23] A. Rao, D. Barkley, G. S. França, and I. Yanai. Exploring tissue architecture using spatial transcriptomics. Nature, 596(7871):211–220, 2021.

[24] D. Righelli, L. M. Weber, H. L. Crowell, B. Pardo, L. Collado-Torres, S. Ghazanfar, A. T. Lun, S. C. Hicks, and D. Risso. SpatialExperiment: infrastructure for spatially-resolved transcriptomics data in R using Bioconductor. Bioinformatics, 38(11):3128–3131, 2022.

[25] M. E. Ritchie, B. Phipson, D. Wu, Y. Hu, C. W. Law, W. Shi, and G. K. Smyth. limma powers differential expression analyses for RNA-sequencing and microarray studies. Nucleic acids research, 43(7):e47–e47, 2015.

[26] M. D. Robinson and A. Oshlack. A scaling normalization method for differential expression analysis of RNA-seq data. Genome biology, 11(3):1–9, 2010.

[27] D. Ruben, Z. Qian, D. Rui, C.-H. L. Eng, L. Huipeng, L. Kan, F. Yuntian, Z. Tianxiao, S. Arpan, B. Feng, R. E. George, P. Nico, C. Long, and Y. Guo-Cheng. Giotto: a toolbox for integrative analysis and visualization of spatial expression data. Genome Biol, 22:78, 2021.

[28] A. G. C. A. M. E. M. C. R. V. J. W. L. M. C. F. C.. E. Z. M. Samuel G Rodriques, Robert R Stickels. Slide-seq: A scalable technology for measuring genome-wide expression at high spatial resolution. Science (New York, N.Y.), 363(6434):1463–1467, 2019.

[29] R. Satija, J. A. Farrell, D. Gennert, A. F. Schier, and A. Regev. Spatial reconstruction of single-cell gene expression data. Nature biotechnology, 33(5):495– 502, 2015.

[30] W. Z. L. C. Sheel Shah, Eric Lubeck. In situ transcription profiling of single cells reveals spatial organization of cells in the mouse hippocampus. Neuron, 92(2):342–357, 2016.

[31] J. Z. Shiquan Sun and X. Zhou. Statistical analysis of spatial expression patterns for spatially resolved transcriptomic studies. Nature Methods, 17:193– 200, 2020.

[32] J. Z. Shiquan Sun and X. Zhou. SPARK-X: non-parametric modeling enables scalable and robust detection of spatial expression patterns for large spatial transcriptomic studies. Genome Biology, 22:184, 2021.

[33] R. R. Stickels, E. Murray, P. Kumar, J. Li, J. L. Marshall, D. J. Di Bella, P. Arlotta, E. Z. Macosko, and F. Chen. Highly sensitive spatial transcriptomics at near-cellular resolution with Slide-seqV2. Nature Biotechnology, 39:313–319, March 2021.

[34] P. L. Ståhl, F. Salmén, S. Vickovic, A. Lundmark, J. F. Navarro, J. Magnusson, S. Giacomello, M. Asp, J. O. Westholm, M. Huss, A. Mollbrink, S. Linnarsson, S. Codeluppi, Åke Borg, F. Pontén, P. I. Costea, P. Sahlén, J. Mulder, O. Bergmann, J. Lundeberg, and J. Frisén. Visualization and analysis of gene expression in tissue sections by spatial transcriptomics. Science, 353(6294):78–82, 2016.

[35] K. Thrane, H. Eriksson, J. Maaskola, J. Hansson, and J. Lundeberg. Spatially resolved transcriptomics enables dissection of genetic heterogeneity in stage iii cutaneous malignant melanoma. Cancer Research, 78(20):5970–5979, 2018.

[36] S. A. T. Valentine Svensson and O. Stegle. SpatialDE: identification of spatially variable genes. Nature Methods, 15:343–346, 2018.

[37] S. Vickovic, G. Eraslan, F. Salmén, J. Klughammer, L. Stenbeck, D. Schapiro, T. Äijö, R. onneau, L. Bergenstråhle, J. F. Navarro, J. Gould, G. K. Griffin, å. Borg, M. Ronaghi, J. Frisén, J. Lundeberg, A. Regev, and P. L. Ståhl. High-definition spatial transcriptomics for in situ tissue profiling. Nature methods, 16:987–990, 2019.

[38] L. M. Weber, A. Saha, A. Datta, K. D. Hansen, and S. C. Hicks. nnSVG: scalable identification of spatially variable genes using nearest-neighbor Gaussian processes. bioRxiv, May 2022.

[39] S. S. Wilks. The large-sample distribution of the likelihood ratio for testing composite hypotheses. The annals of mathematical statistics, 9(1):60–62, 1938.

[40] E. Zhao, M. R. Stone, X. Ren, J. Guenthoer, K. S. Smythe, T. Pulliam, S. R. Williams, C. R. Uytingco, S. E. Taylor, P. Nghiem, et al. Spatial transcriptomics at subspot resolution with BayesSpace. Nature Biotechnology, 39(11):1375–1384, 2021.

[41] P. Zhao, J. Zhu, Y. Ma, and X. Zhou. Modeling zero inflation is not necessary for spatial transcriptomics. Genome Biology, 23(1):118, 2022.

