## Supplementary material for "*DESpace*: spatially variable gene detection via differential expression testing of spatial clusters"


#### 1 Supplementary Details

##### 1.1 Adjusted p-values

In all methods, to control for the false discovery rate, we used adjusted p-values, obtained via Benjamini-Hochberg correction [1].

##### 1.2 Quality control and filtering

Quality control followed the guidelines in the online book entitled "Orchestrating Spatially-Resolved Transcriptomics Analysis with Bioconductor" [3]. In particular, low quality spots were identified using *addPerCellQC* function from *Scuttle* R package [2]. Based on total UMI counts per spot, number of genes with non-zero UMI counts per spot (detected genes), proportion of read mapping to mitochondrial genes, and number of cells per spot, low quality spots were filtered with sample-specific thresholds. Furthermore, undetected genes and lowly expressed genes were removed to ensure that each gene had at least 20 non-zero spots of expression.

##### 1.3 Simulating SV patterns

To generate our simulations, we initially partitioned the tissue into two regions, $R_1$ and $R_2$, with $n_1$ and $n_2$ spots, respectively; as described above, we generated a variety of spatial structures. Then, starting from these partitions, we took advantage of the *runPatternSimulation* function from *Giotto*'s R package [5]. In particular, *runPatternSimulation* considers the most highly abundant $n_1$ genes and assigns them to region $R_1$ with probability $\pi$. When $\pi = 1$, all the most highly abundant $n_1$ spots are assigned to region $R_1$, while $\pi = 0.5$ leads to uniform patterns.

Our simulations are composed of a mixture of uniform and SV genes: we obtained uniform patterns by setting $\pi = 0.5$, while SVGs were simulated using $\pi = 0.9$. Only in the multi-sample simulation, SVGs were simulated using $\pi = 0.7$; we choose to decrease $\pi$ because the TPRs of both single-sample and multi-sample modes were close to 1: decreasing $\pi$ allowed us to show a difference between the TPRs of the two modes. For the strong *vs.* weak patterns simulation, we set *pi* to 0.9 and 0.6 for the former and latter case, respectively.

In the *bottom/right*, *circular*, and *annotations* patterns, we simulated 1/3 of SVGs and 2/3 of uniform genes; in each one of these simulations, half of the SVGs are highly abundant one region (e.g., centre of the circle, or Layer 5; see Figure 2), and half are highly abundant in the complementary region (e.g., outside the circle, or outside Layer 5; see Figure 2). In *mixture* and *inverted mixture* simulations, we generate 50% of uniform genes and 50% of SVGs, in equal proportions from: 5, 3 and 4 distinct patterns in the *LIBD*, *melanoma*, and *mouse cerebellum* datasets, respectively.

##### 1.4 Individual cluster testing

Computing dispersion estimates is the most computationally demanding task in *edgeR*. Since this step depends on the design matrix of the test performed, for each cluster being tested, *edgeR* requires re-computing such dispersion estimates. In order to accelerate calculations, we also provide an alternative implementation, where we use the dispersion estimates previously computed for the gene-level test. This results in a significant decrease in the computational burden (runtime decreased, on average, from 32 to 2 minutes per sample), resulting in analogous performance (Table 1, and Supplementary Table 2).

##### 1.5 Software versions

Most analyses were performed in R (version 4.2.0, with Bioconductor packages from release 3.15); some methods (i.e., *StLearn*, *SpatialDE*, *SpatialDE2* and *SpaGCN*) were run in python (version 3.10.0).

#### 2 Supplementary Tables

| Data set | Sample Ids | Before Filtering | | After Filtering | |
|---|---|---|---|---|---|
| | | # of Genes | # of Spots | # of Genes | # of Spots |
| LIBD | 151507 | 33538 | 4226 | 14226 | 4172 |
| | 151508 | 33538 | 4384 | 13835 | 4333 |
| | 151509 | 33538 | 4789 | 14561 | 4738 |
| | 151510 | 33538 | 4634 | 14331 | 4606 |
| | 151669 | 33538 | 3661 | 14547 | 3647 |
| | 151670 | 33538 | 3498 | 14183 | 3473 |
| | 151671 | 33538 | 4110 | 14926 | 4079 |
| | 151672 | 33538 | 4015 | 14634 | 3989 |
| | 151673 | 33538 | 3639 | 15115 | 3604 |
| | 151674 | 33538 | 3673 | 15878 | 3634 |
| | 151675 | 33538 | 3592 | 14626 | 3556 |
| | 151676 | 33538 | 3460 | 14672 | 3405 |
| | Average | 33538 | 3973 | 14628 | 3936 |
| melanoma | mel1-rep1 | 15666 | 280 | 7962 | 276 |
| | mel1_rep2 | 16148 | 294 | 9104 | 292 |
| | mel2_rep1 | 16831 | 384 | 9315 | 380 |
| | mel2_rep2 | 16605 | 381 | 8220 | 375 |
| | mel3_rep1 | 15653 | 257 | 7604 | 253 |
| | mel3_rep2 | 15769 | 295 | 7882 | 290 |
| | mel4_rep1 | 14408 | 213 | 5944 | 208 |
| | mel4_rep2 | 15991 | 249 | 9351 | 246 |
| | Average | 15884 | 294 | 8173 | 290 |
| mouse cerebellum | | 20141 | 11626 | 8277 | 11615 |

**Supplementary Table 1:** Number of genes and spots for each sample, before and after filtering.

| Pattern | Sample | *DESpace* | | *FindAllMarkers* | | *findMarkers* | |
|---|---|---|---|---|---|---|---|
| | | *BayesSpace* | *StLearn* | *BayesSpace* | *StLearn* | *BayesSpace* | *StLearn* |
| Mixture | 151507 | 99.6 | 98.7 | 99.5 | 99.3 | 98.6 | 96.9 |
| | 151669 | 99.4 | 99.5 | 99.8 | 99.8 | 97.7 | 97.8 |
| | 151673 | 99.8 | 99.0 | 99.8 | 99.6 | 99.1 | 98.7 |
| | Average | 99.6 | 99.1 | 99.6 | 99.6 | 98.5 | 97.8 |
| Inverted Mixture | 151507 | 86.6 | 79.7 | 81.1 | 77.7 | 36.1 | 73.8 |
| | 151669 | 89.3 | 88.8 | 84.2 | 85.4 | 59.9 | 83.3 |
| | 151673 | 90.9 | 85. | 87.4 | 84.5 | 86.0 | 82.4 |
| | Average | 88.9 | 84.7 | 84.2 | 82.5 | 60.7 | 79.8 |

**Supplementary Table 2:** Individual cluster results, for each sample. Percentage of times that *DESpace*, *Seurat*'s *FindAllMarkers* and *scran*'s *findMarkers* identified the main SV cluster in *mixture* (i.e., Mixture) and *inverted mixture* simulations (i.e., Inverted), using *BayesSpace* and *StLearn* clusters.

| Pattern | Sample | *BayesSpace* | *StLearn* |
|---|---|---|---|
| Mixture | 151507 | 99.6 | 98.7 |
| | 151669 | 99.5 | 99.5 |
| | 151673 | 99.8 | 99.0 |
| | Average | 99.6 | 99.1 |
| Inverted Mixture | 151507 | 86.7 | 79.9 |
| | 151669 | 89.3 | 88.7 |
| | 151673 | 90.8 | 85.5 |
| | Average | 88.9 | 84.7 |

**Supplementary Table 3:** Individual cluster results. *DESpace* was run without recomputing the dispersion estimates for each spatial cluster tested (i.e., faster mode). Percentage of times that *DESpace* identified the main SV cluster in each *mixture* and *inverted mixture* simulations, using *BayesSpace* or *StLearn* clusters.

| Method \ Pattern | Bottom/Right | Circular | Annotations | Mixture | Inverted mixture | Average |
|---|---|---|---|---|---|---|
| StLearn_findMarkers | 0.96 | 0.98 | 0.97 | 0.74 | 0.64 | 0.86 |
| BayesSpace_DESpace | 0.97 | 0.95 | 0.95 | 0.7 | 0.61 | 0.84 |
| StLearn_DESpace | 0.97 | 0.95 | 0.96 | 0.69 | 0.57 | 0.83 |
| SPARK | 0.96 | 0.95 | 0.92 | 0.69 | 0.56 | 0.82 |
| BayesSpace_FindAllMarkers | 0.94 | 0.91 | 0.91 | 0.72 | 0.57 | 0.81 |
| StLearn_FindAllMarkers | 0.94 | 0.91 | 0.92 | 0.7 | 0.56 | 0.8 |
| SpatialDE2 | 0.96 | 0.93 | 0.85 | 0.66 | 0.55 | 0.79 |
| SpatialDE | 0.93 | 0.89 | 0.84 | 0.71 | 0.56 | 0.79 |
| nnSVG | 0.89 | 0.87 | 0.87 | 0.71 | 0.58 | 0.78 |
| SPARK-X | 0.93 | 0.9 | 0.77 | 0.69 | 0.54 | 0.77 |
| MERINGUE | 0.84 | 0.83 | 0.83 | 0.72 | 0.58 | 0.76 |
| BayesSpace_findMarkers | 0.96 | 0.98 | 0.31 | 0.7 | 0.61 | 0.71 |
| SpaGCN | 0.8 | 0.85 | 0.83 | 0.5 | 0.47 | 0.69 |

**Supplementary Table 4:** Area under the curve values, for the curves in Supplementary Figure 5, left column (weak vs. strong spatial simulations based on *LIBD* data). Rows and columns refer to the method and pattern, respectively.

| Method \ Pattern | Bottom/Right | Circular | Annotations | Mixture | Inverted mixture | Average |
|---|---|---|---|---|---|---|
| BayesSpace_DESpace | 0.99 | 0.97 | 0.99 | 0.79 | 0.76 | 0.9 |
| StLearn_DESpace | 0.99 | 0.97 | 0.99 | 0.78 | 0.75 | 0.9 |
| BayesSpace_FindAllMarkers | 0.98 | 0.95 | 0.98 | 0.78 | 0.76 | 0.89 |
| StLearn_FindAllMarkers | 0.98 | 0.95 | 0.98 | 0.78 | 0.76 | 0.89 |
| BayesSpace_findMarkers | 0.99 | 0.99 | 0.99 | 0.74 | 0.69 | 0.88 |
| StLearn_findMarkers | 0.99 | 0.99 | 0.99 | 0.73 | 0.68 | 0.88 |
| SpaGCN | 0.88 | 0.91 | 0.88 | 0.66 | 0.68 | 0.8 |
| SPARK | 0.95 | 0.88 | 0.83 | 0.7 | 0.65 | 0.8 |
| nnSVG | 0.92 | 0.85 | 0.85 | 0.71 | 0.65 | 0.8 |
| SpatialDE | 0.93 | 0.86 | 0.8 | 0.71 | 0.65 | 0.79 |
| SpatialDE2 | 0.91 | 0.85 | 0.8 | 0.69 | 0.63 | 0.78 |
| SPARK-X | 0.9 | 0.87 | 0.76 | 0.66 | 0.6 | 0.76 |
| MERINGUE | 0.86 | 0.8 | 0.8 | 0.68 | 0.61 | 0.75 |

**Supplementary Table 5:** Area under the curve values, for the curves in Supplementary Figure 5, right column (weak vs. strong spatial simulations based on *melanoma* data). Rows and columns refer to the method and pattern, respectively.

| Method \ Keyword | HLA | melanoma marker | melanoma | overall |
|---|---|---|---|---|
| BayesSpace_DESpace | 65 | 8 | 1073 | 1107 |
| StLearn_DESpace | 58 | 7 | 1062 | 1094 |
| BayesSpace_FindAllMarkers | 40 | 7 | 1069 | 1092 |
| StLearn_FindAllMarkers | 50 | 5 | 1049 | 1078 |
| SPARK-X | 48 | 7 | 1042 | 1068 |
| StLearn_findMarkers | 57 | 6 | 1035 | 1064 |
| nnSVG | 64 | 6 | 1021 | 1055 |
| SPARK | 63 | 7 | 1011 | 1044 |
| SpatialDE2 | 50 | 5 | 1006 | 1030 |
| MERINGUE | 64 | 8 | 995 | 1029 |
| SpatialDE | 56 | 7 | 984 | 1014 |
| BayesSpace_findMarkers | 50 | 6 | 961 | 986 |
| SpaGCN | 36 | 5 | 913 | 936 |

**Supplementary Table 6:** Overall number of genes detected, among the top 500 results returned by each method, on every sample from the *melanoma* dataset. The lists of interesting genes, were found by searching on The Human Protein Atlas website for the following keywords: "HLA", "melanoma marker" and "melanoma" (HLA results were filtered to include HLA genes only). "overall" represents an aggregation of the three lists. Globally, "HLA", "melanoma marker", "melanoma", and "overall" contain a total of 19, 12, 3,398, and 3,410 genes, respectively.

#### 3 Supplementary Figures

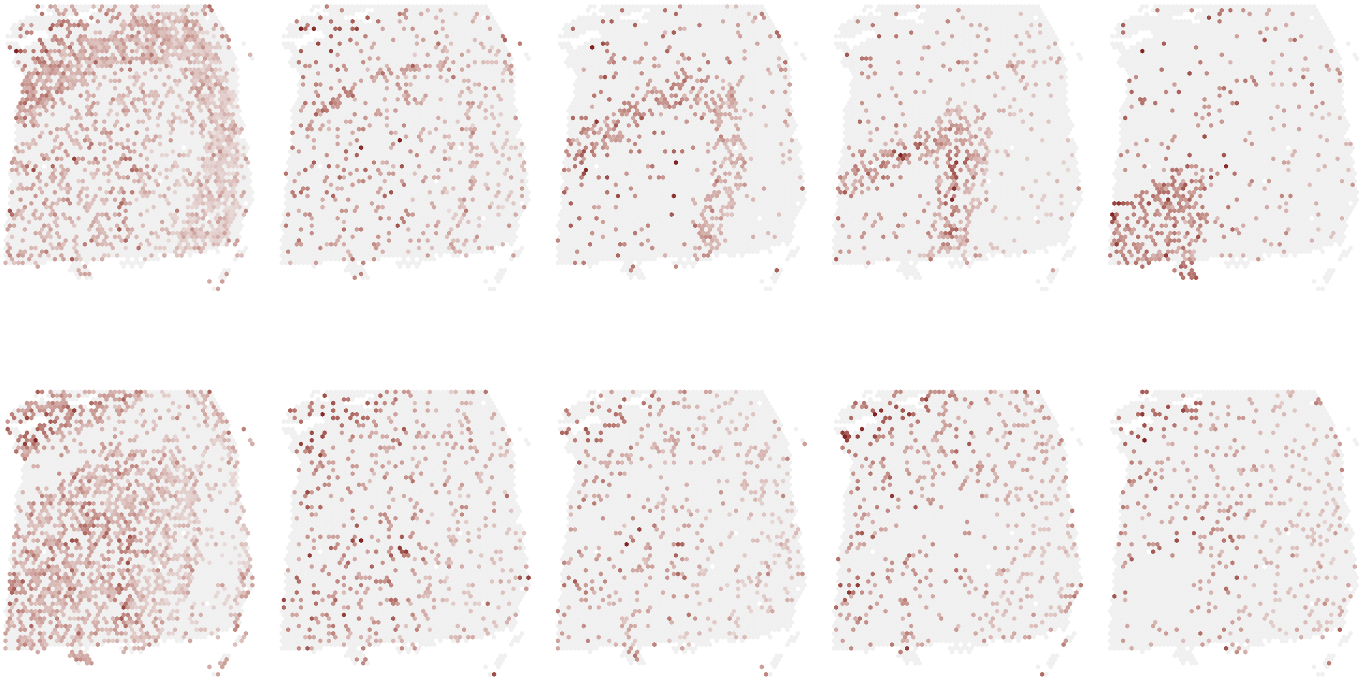

**Supplementary Figure 1:** Examples of simulates SVGs, from the *LIBD* data, following *mixture* (top row) and *inverted mixture* (bottom row) SV patterns. From left to right, in the top (bottom) row, the highly (lowly) abundant regions refer to layer 3, layer 4, layer 5, layer 6, and white matter.

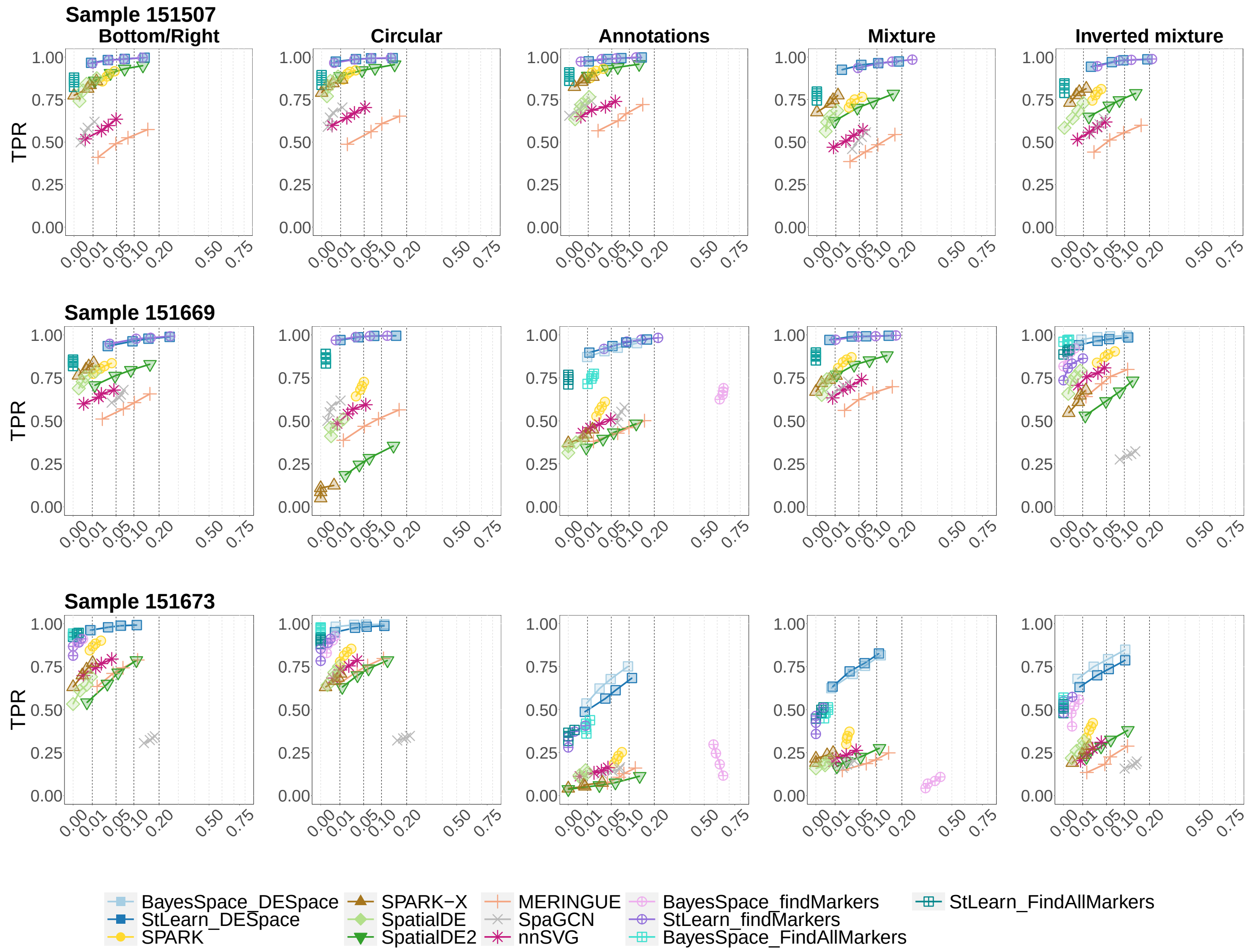


**Supplementary Figure 2:** TPR vs. FDR for SVG detections in the individual sample simulations, in each *LIBD* sample. Rows and columns refer to the specific sample used as anchor data in the simulation, and to the SV profiles, respectively. *BayesSpace_ DESpace* and *StLearn_ DESpace* indicate *DESpace* based on spatial clusters computed via *BayesSpace* and *StLearn*, respectively.

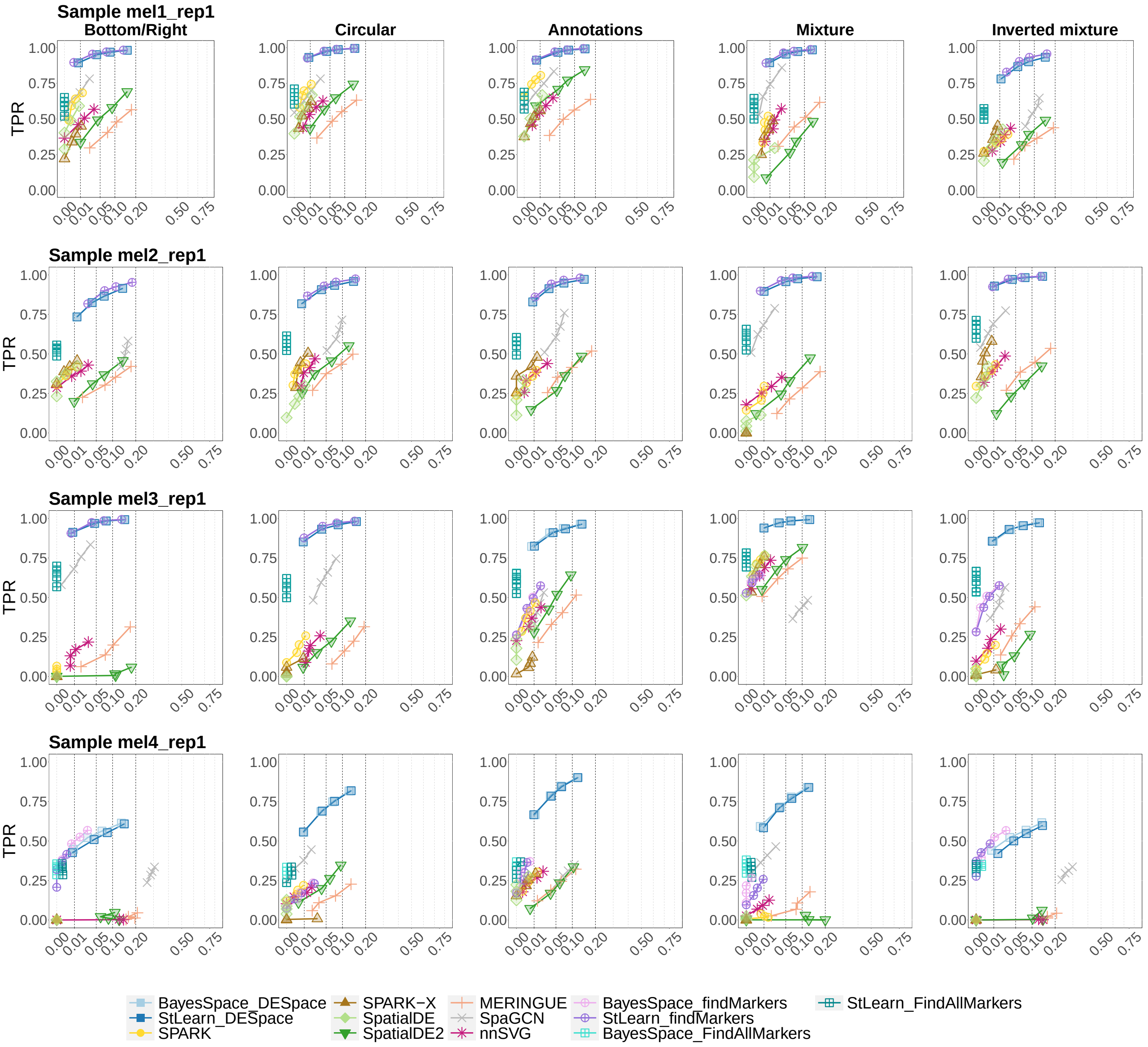


**Supplementary Figure 3:** TPR vs. FDR for SVG detections in the individual sample simulations, in each *melanoma* sample. Rows and columns refer to the specific sample used as anchor data in the simulation, and to the SV profiles, respectively. *BayesSpace_ DESpace* and *StLearn_ DESpace* indicate *DESpace* based on spatial clusters computed via *BayesSpace* and *StLearn*, respectively.

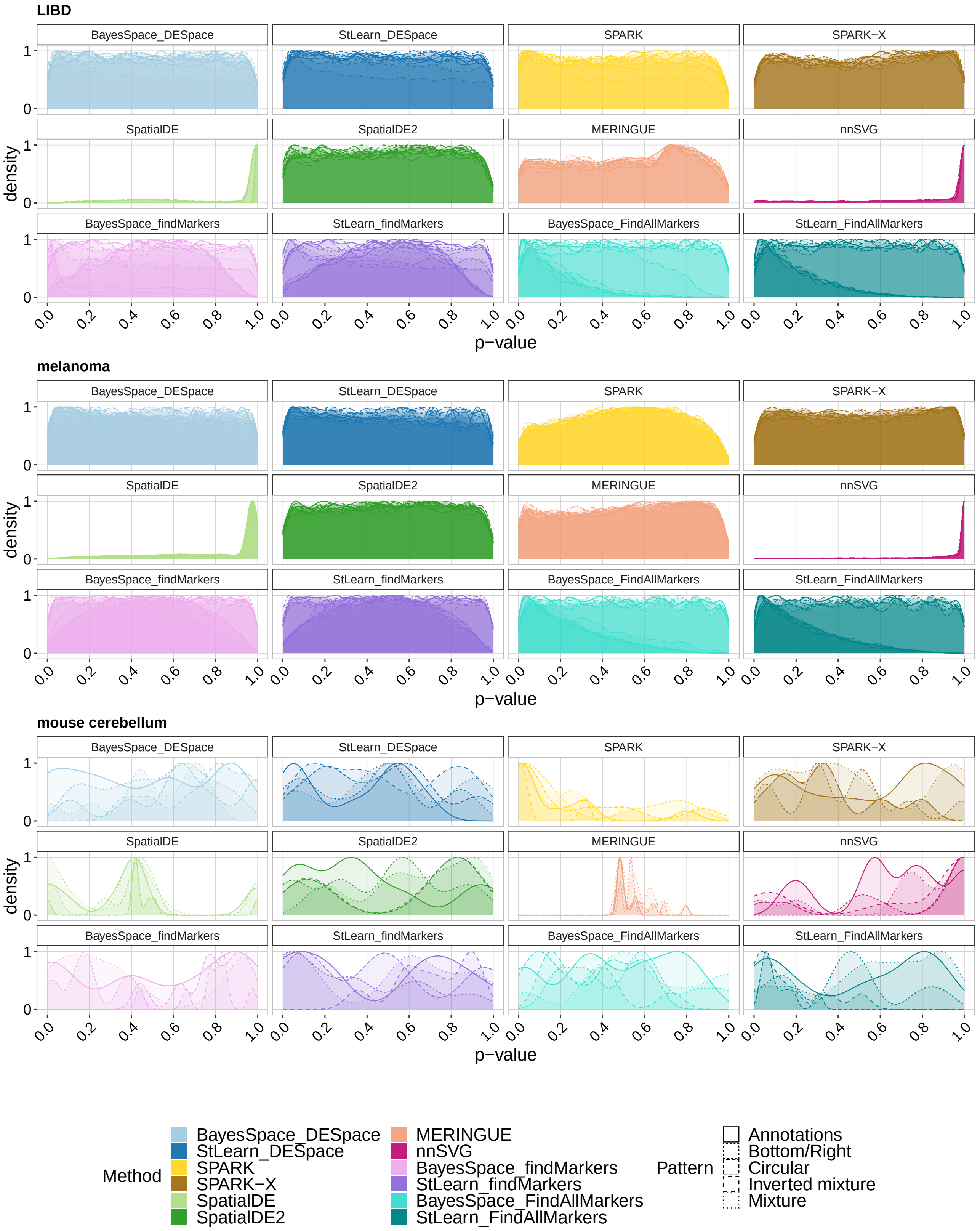


**Supplementary Figure 4:** Density of raw p-values in the null individual sample simulations (i.e., no differences between groups), based on *LIBD*, *melanoma*, and *mouse cerebellum* anchor dataset. Each line represents a different simulation, based on a distinct replicate and spatial pattern. *BayesSpace_ DESpace*, *BayesSpace_ findMarkers*, and *BayesSpace_ FindAllMarkers*, as well as their counterparts *StLearn_ DESpace*, *StLearn_ findMarkers*, and *StLearn_ FindAllMarkers*, indicate *DESpace*, *scran*'s *findMarkers*, and *Seurat*'s *FindAllMarkers*, respectively, based on spatial clusters computed via *BayesSpace* and *StLearn*. Note that *SpaGCN* was not included in the analysis, because it does not provide raw p-values.

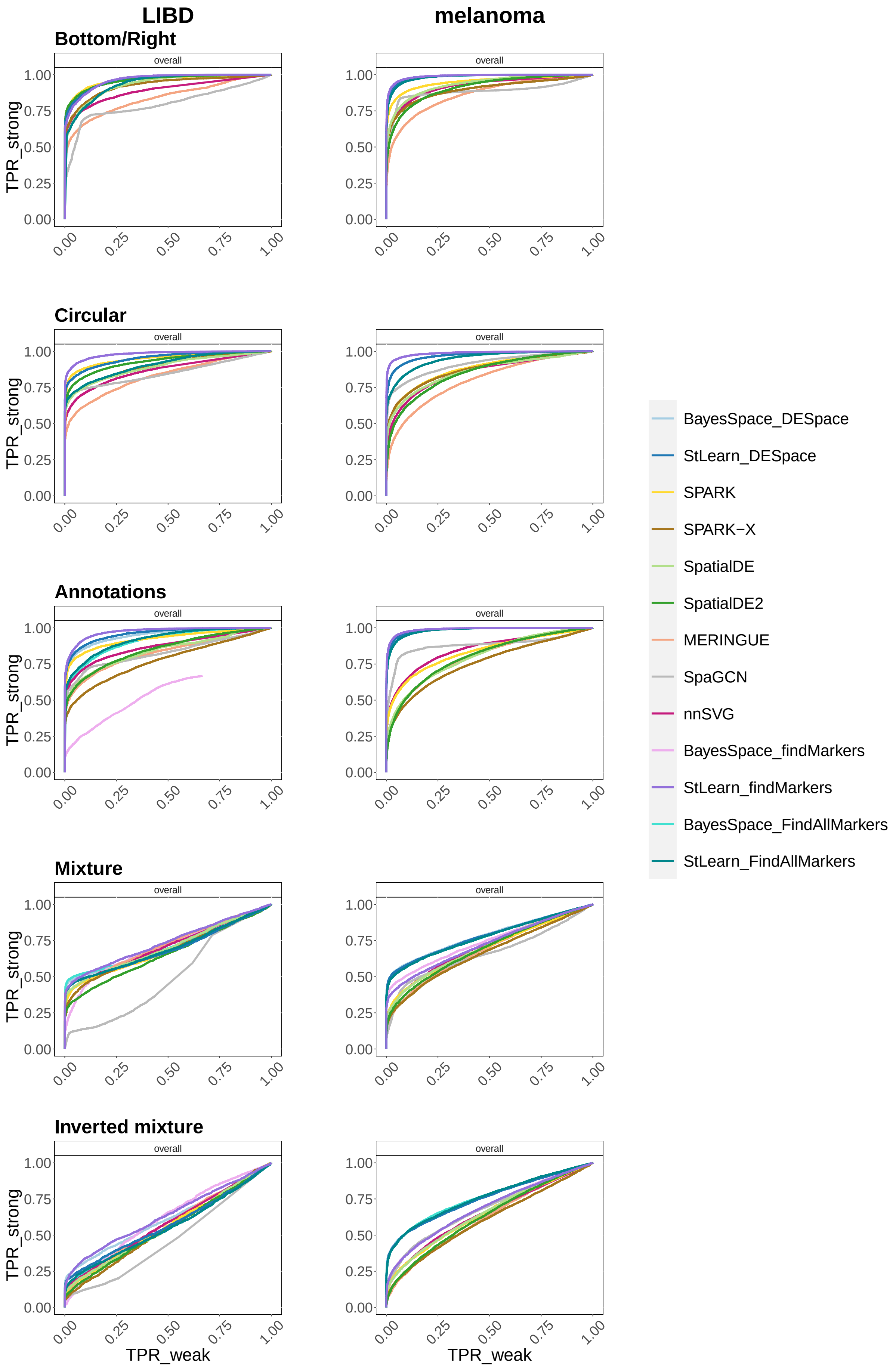


**Supplementary Figure 5:** TPRs for SVGs with weak vs. strong spatial patterns. Rows and columns refer to the SV profile, and the anchor dataset, respectively. *BayesSpace_ DESpace* and *StLearn_ DESpace* indicate *DESpace* based on spatial clusters computed via *BayesSpace* and *StLearn*, respectively.

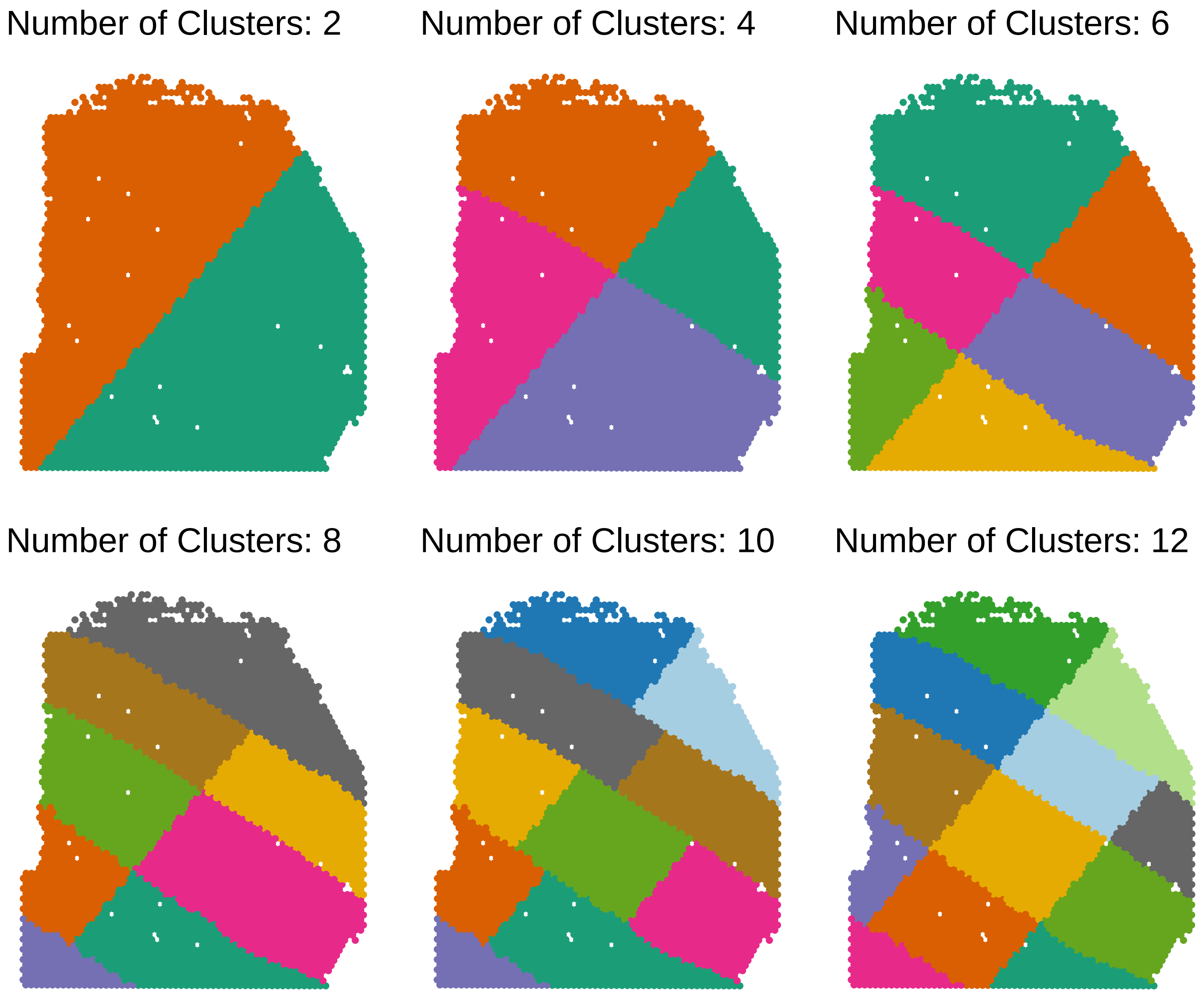


**Supplementary Figure 6:** Artificially generated clusters, based on *LIBD* sample 151507, in the simulation when varying the number of clusters from 2 to 12.

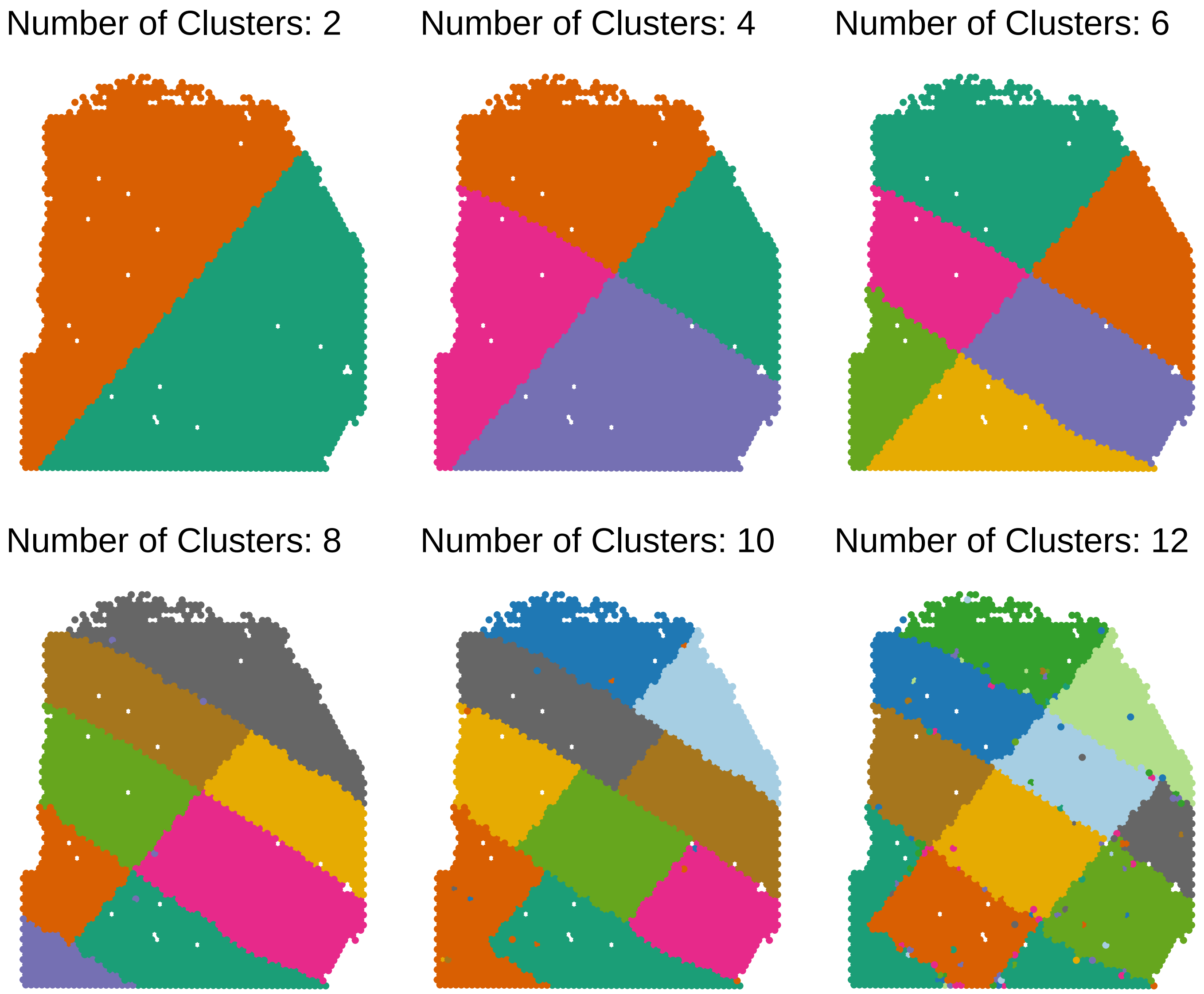


**Supplementary Figure 7:** Spatial clusters inferred by *BayesSpace* in the simulation when varying the number of clusters from 2 to 12, based on *LIBD* sample 151507. Original clusters are shown in Supplementary Figure 6.

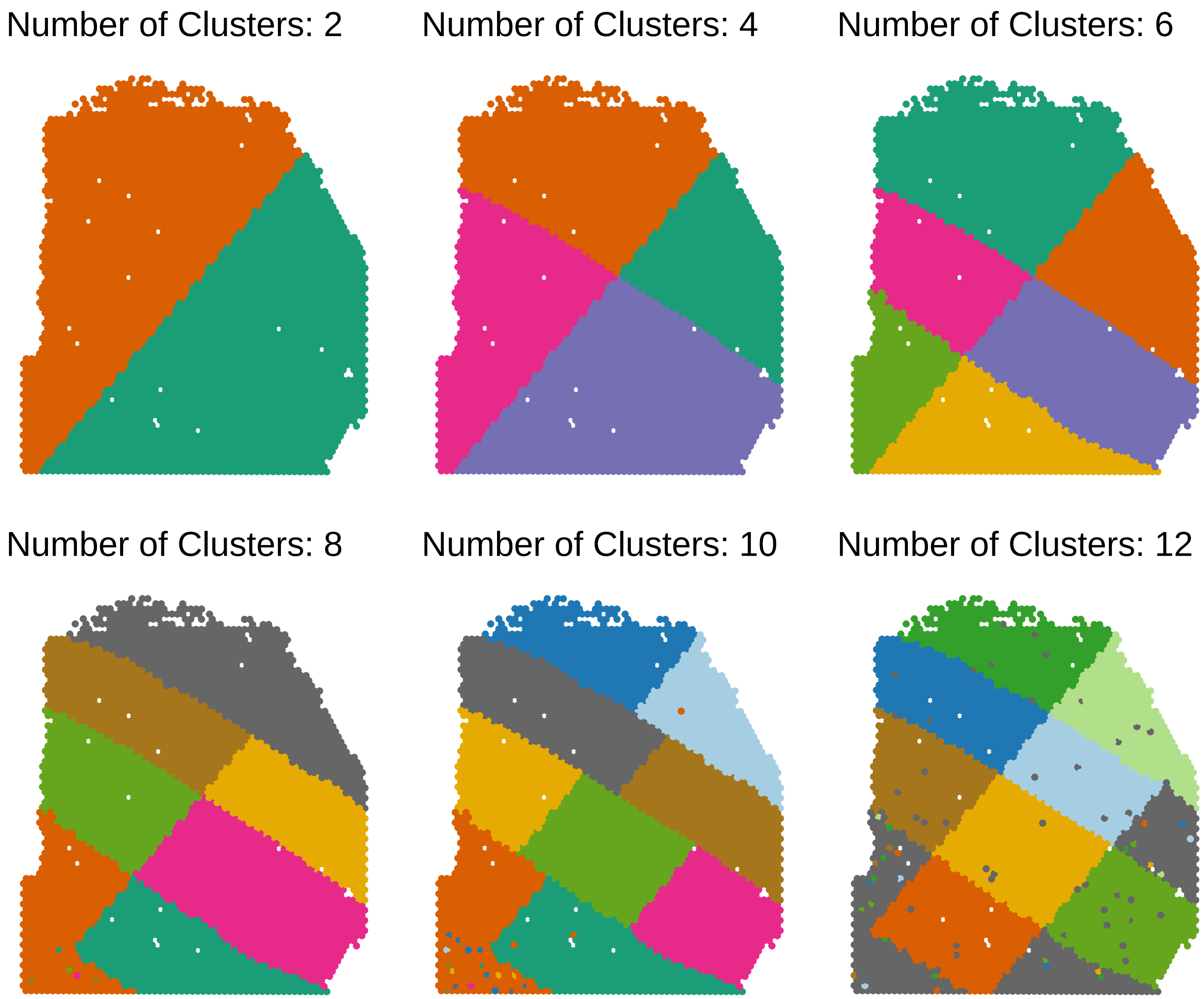


**Supplementary Figure 8:** Spatial clusters inferred by *StLearn* in the simulation when varying the number of clusters from 2 to 12, based on *LIBD* sample 151507. Original clusters are shown in Supplementary Figure 6.

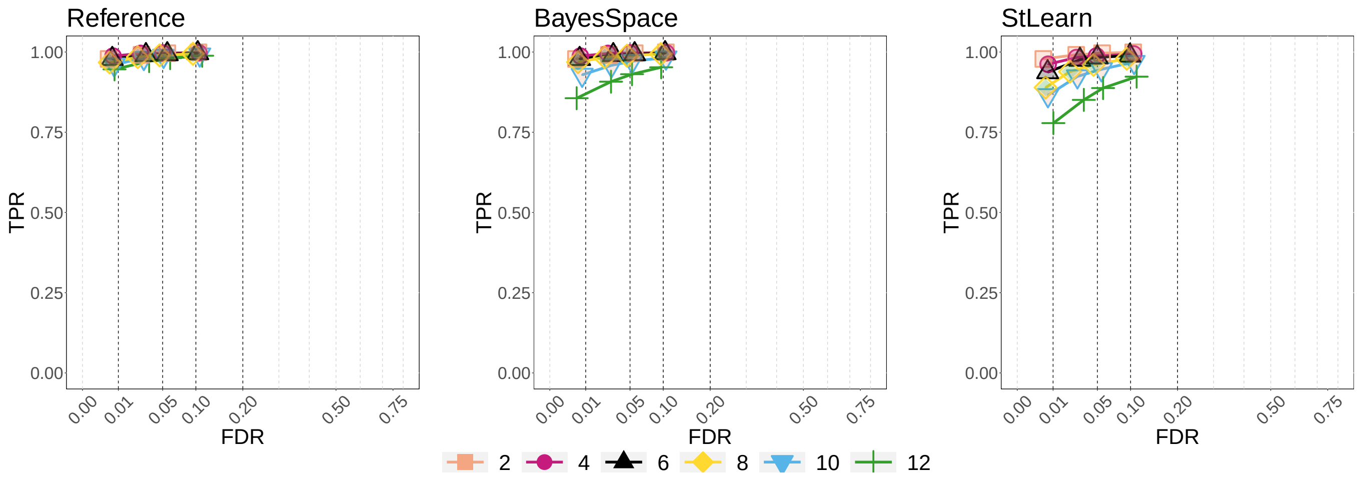


**Supplementary Figure 9:** TPR vs. FDR for SVG detections, based on *LIBD* sample 151507, in the simulation when varying the number of clusters from 2 to 12. *DESpace* uses: i) the original clusters ("Reference") the data was simulated from, shown in Supplementary Figure 6 (left panel); ii) spatial clusters estimated via *BayesSpace*, shown in Supplementary Figure 7 (middle panel); ii) spatial clusters estimated via *StLearn*, shown in Supplementary Figure 8 (right panel).

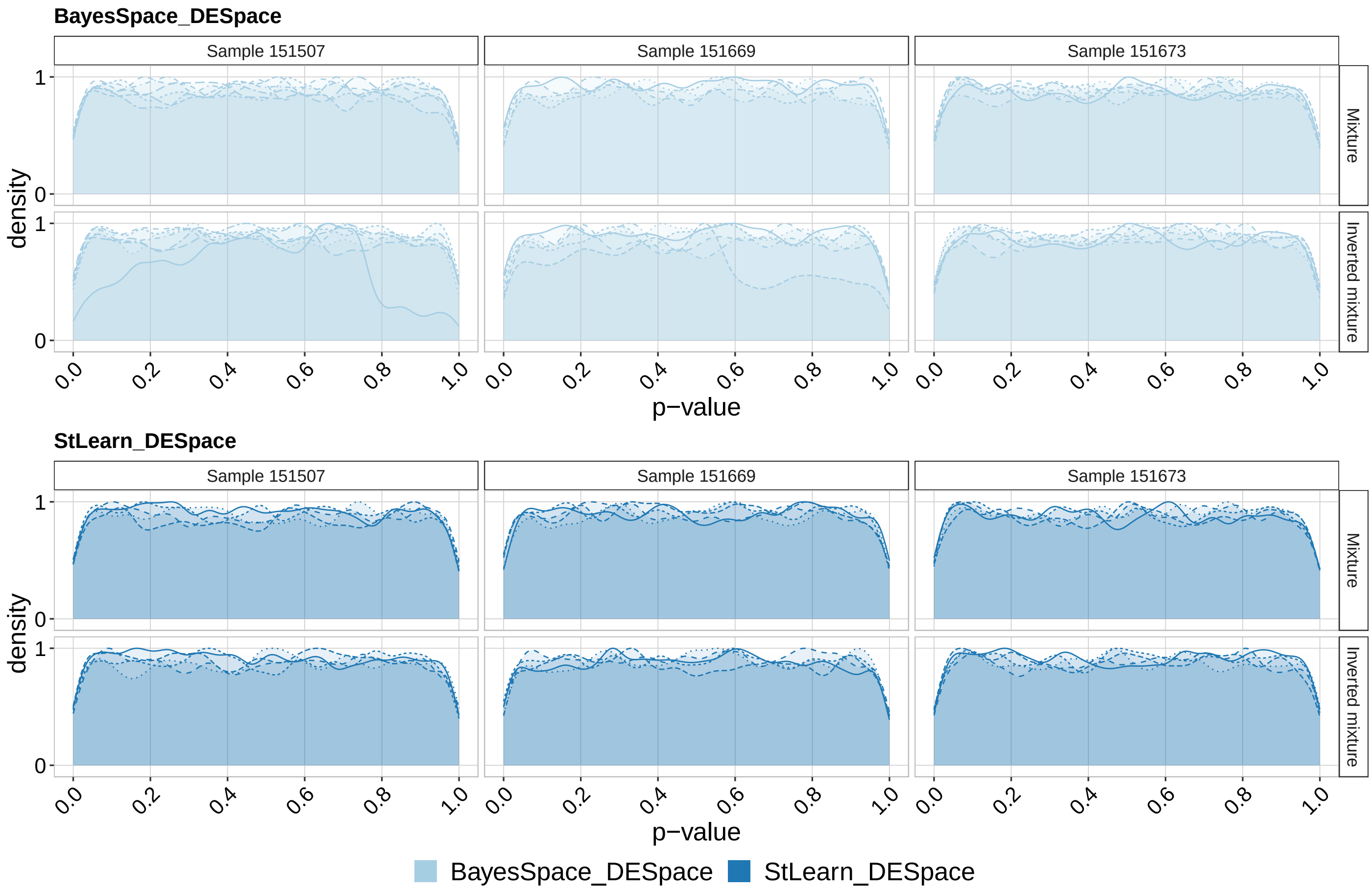


**Supplementary Figure 10:** Density of raw p-values in the null individual cluster simulations (i.e., no differences between groups), based on *LIBD* data. *DESpace* was run recomputing the dispersion estimates for each spatial cluster tested (i.e., slower mode). Each line represents a different simulation, based on a distinct replicate. *BayesSpace_ DESpace* and *StLearn_ DESpace* indicate *DESpace* based on spatial clusters computed via *BayesSpace* and *StLearn*, respectively.

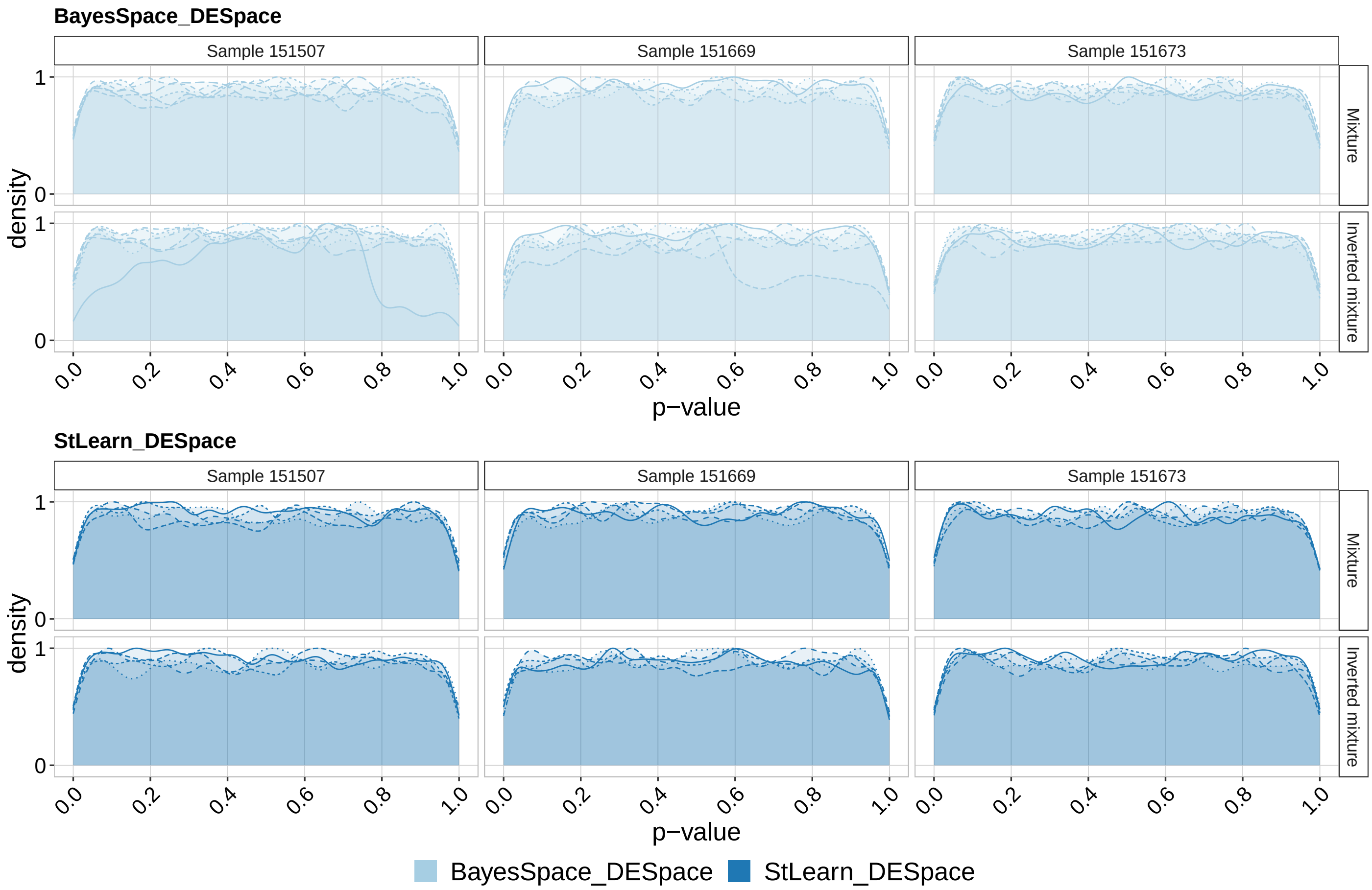


**Supplementary Figure 11:** Density of raw p-values in the null individual cluster simulations (i.e., no differences between groups), based on *LIBD* data. *DESpace* was run without recomputing the dispersion estimates for each spatial cluster tested (i.e., faster mode). Each line represents a different simulation, based on a distinct replicate. *BayesSpace_ DESpace* and *StLearn_ DESpace* indicate *DESpace* based on spatial clusters computed via *BayesSpace* and *StLearn*, respectively.

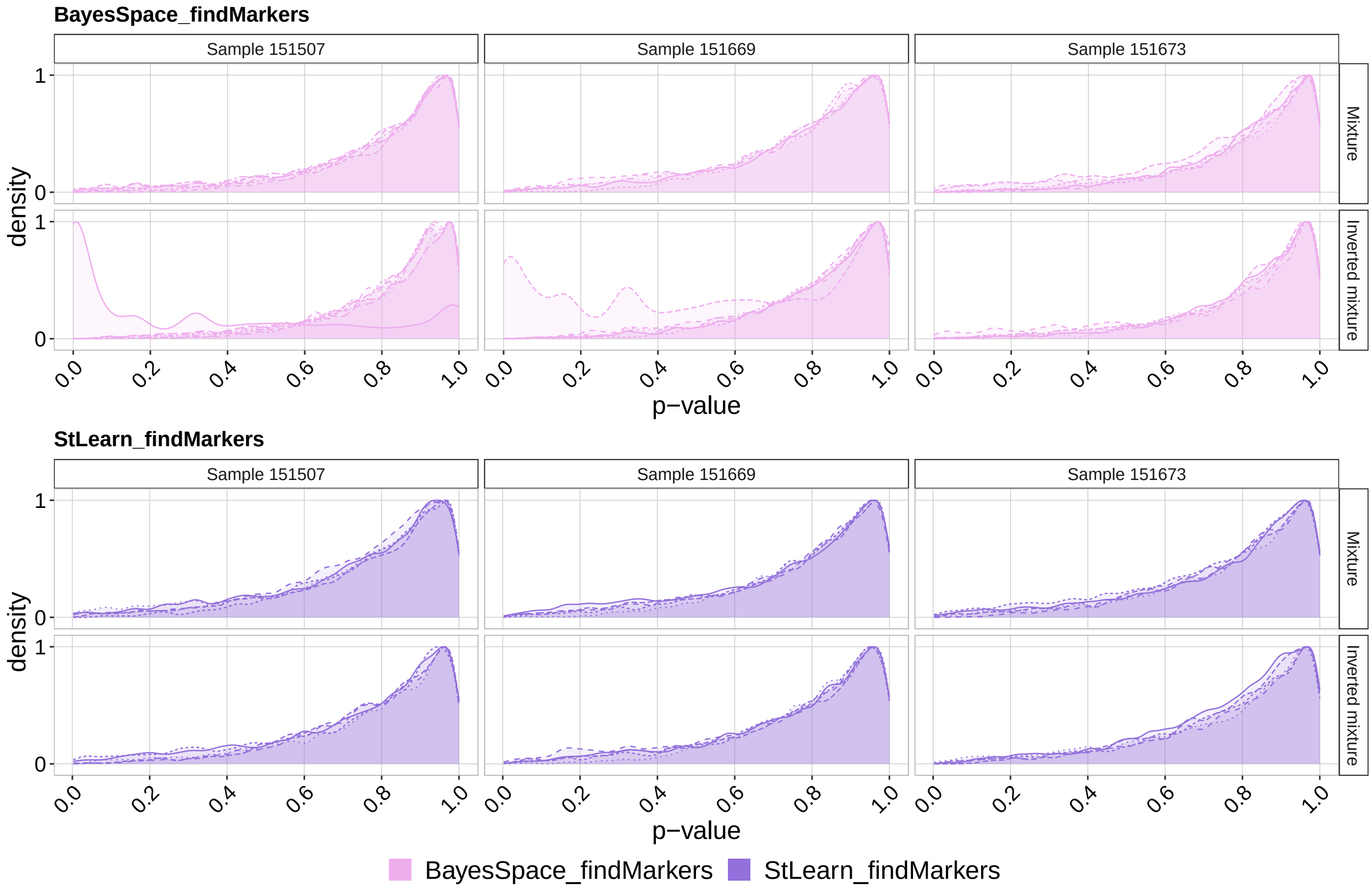


**Supplementary Figure 12:** Density of raw p-values in the null individual cluster simulations (i.e., no differences between groups), based on *LIBD* data. Each line represents a different simulation, based on a distinct replicate. *BayesSpace_ findMarkers* and *StLearn_ findMarkers* indicate *scran*'s *findMarkers* [4] based on spatial clusters computed via *BayesSpace* and *StLearn*, respectively.

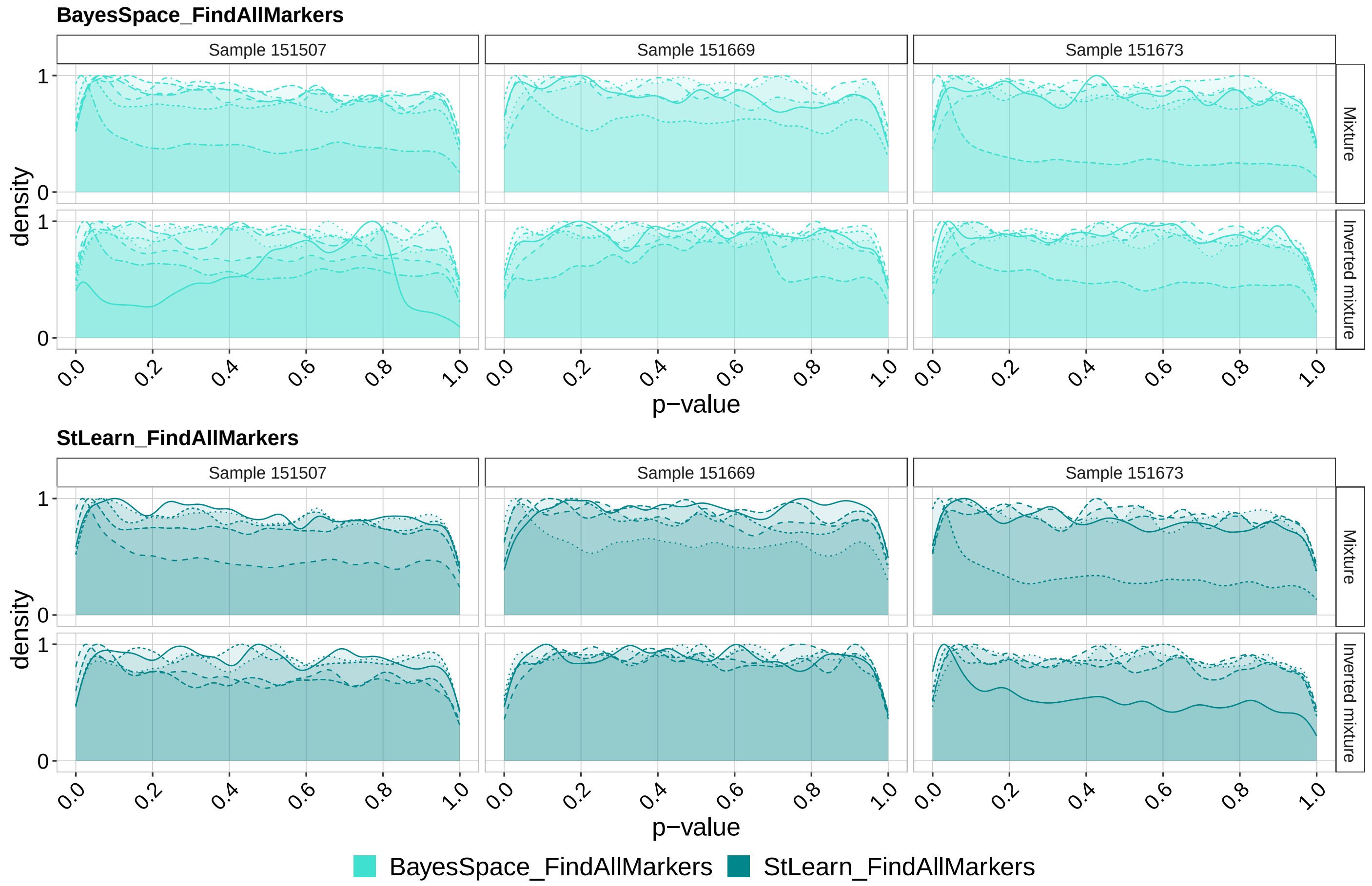


**Supplementary Figure 13:** Density of raw p-values in the null individual cluster simulations (i.e., no differences between groups), based on *LIBD* data. Each line represents a different simulation, based on a distinct replicate. *BayesSpace_ FindAllMarkers* and *StLearn_ FindAllMarkers* indicate *Seurat*'s *FindAllMarkers* [6] based on spatial clusters computed via *BayesSpace* and *StLearn*, respectively.

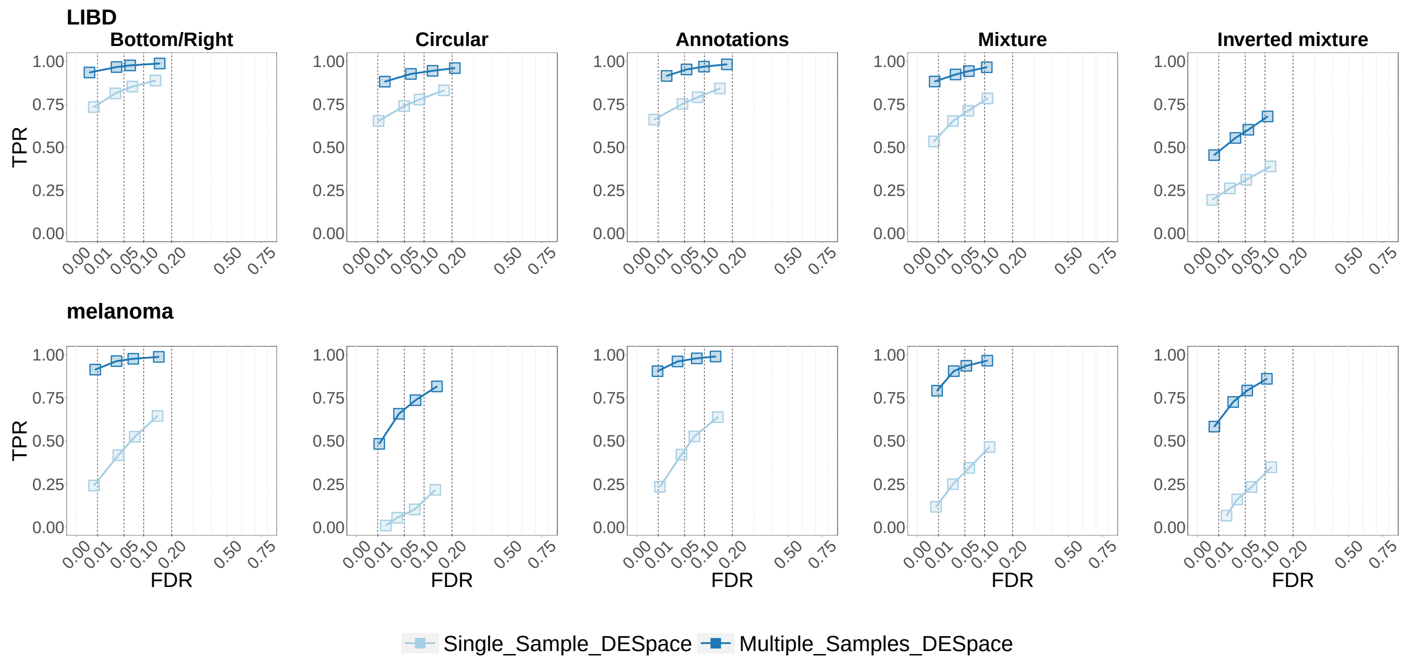


**Supplementary Figure 14:** TPR vs. FDR for SVG detections in the multiple sample simulation. Rows and columns refer to the anchor data used in the simulation, and to the SV profiles, respectively. *Single_ sample_ DESpace* and *Multiple_ samples_ DESpace* indicate *DESpace* on single-sample and multi-sample modes, respectively. Note that TPRs are lower than in the individual simulation, because we have simulated slightly weaker spatial patterns here (see Supplementary Details).

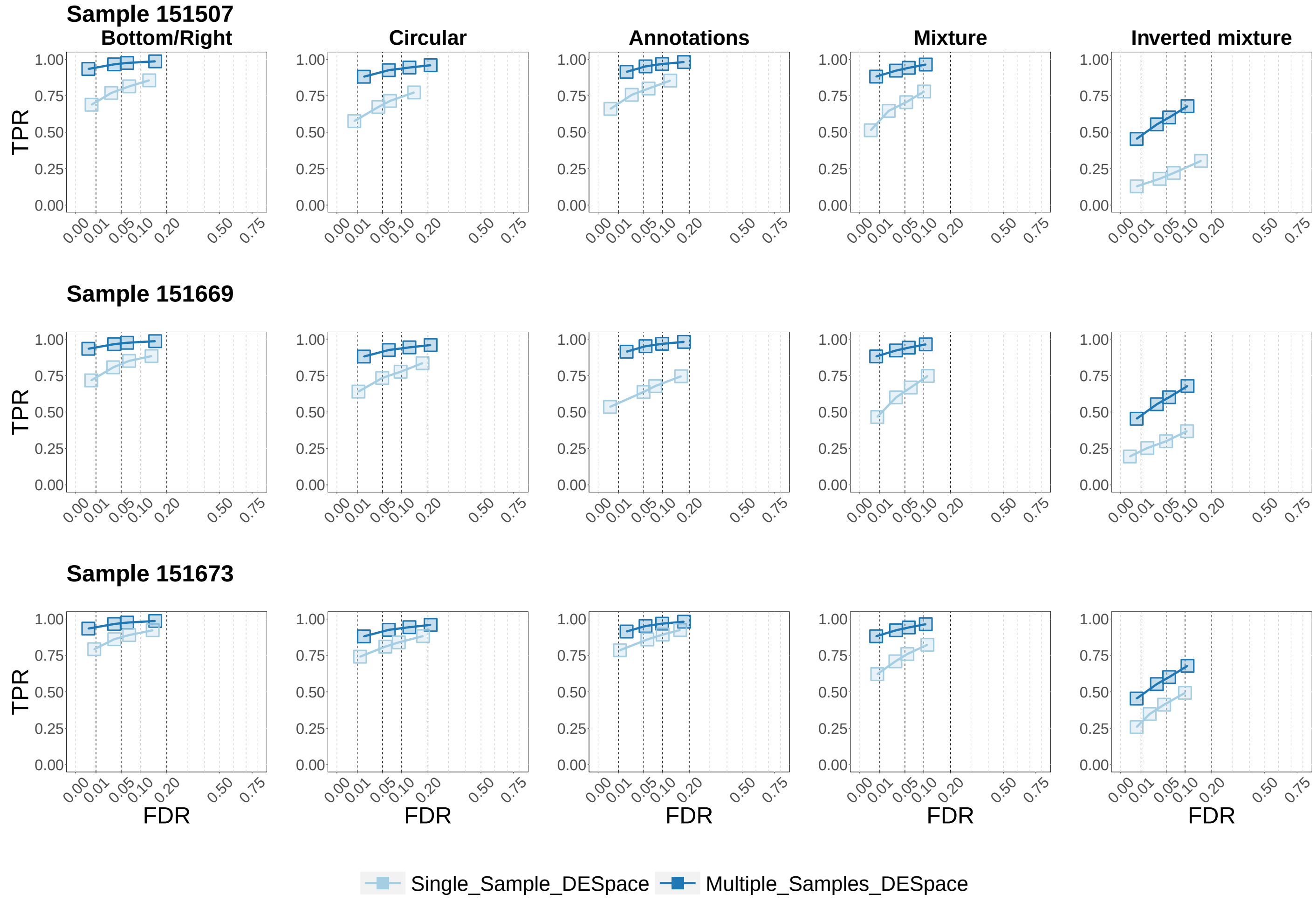


**Supplementary Figure 15:** TPR vs. FDR for SVG detections in the multi-sample simulations, in each *LIBD* sample. Rows and columns refer to the specific sample used as anchor data in the simulation, and to the SV profiles, respectively. *Single_ sample_ DESpace* and *Multiple_ samples_ DESpace* indicate *DESpace* on single-sample and multi-sample modes, respectively.

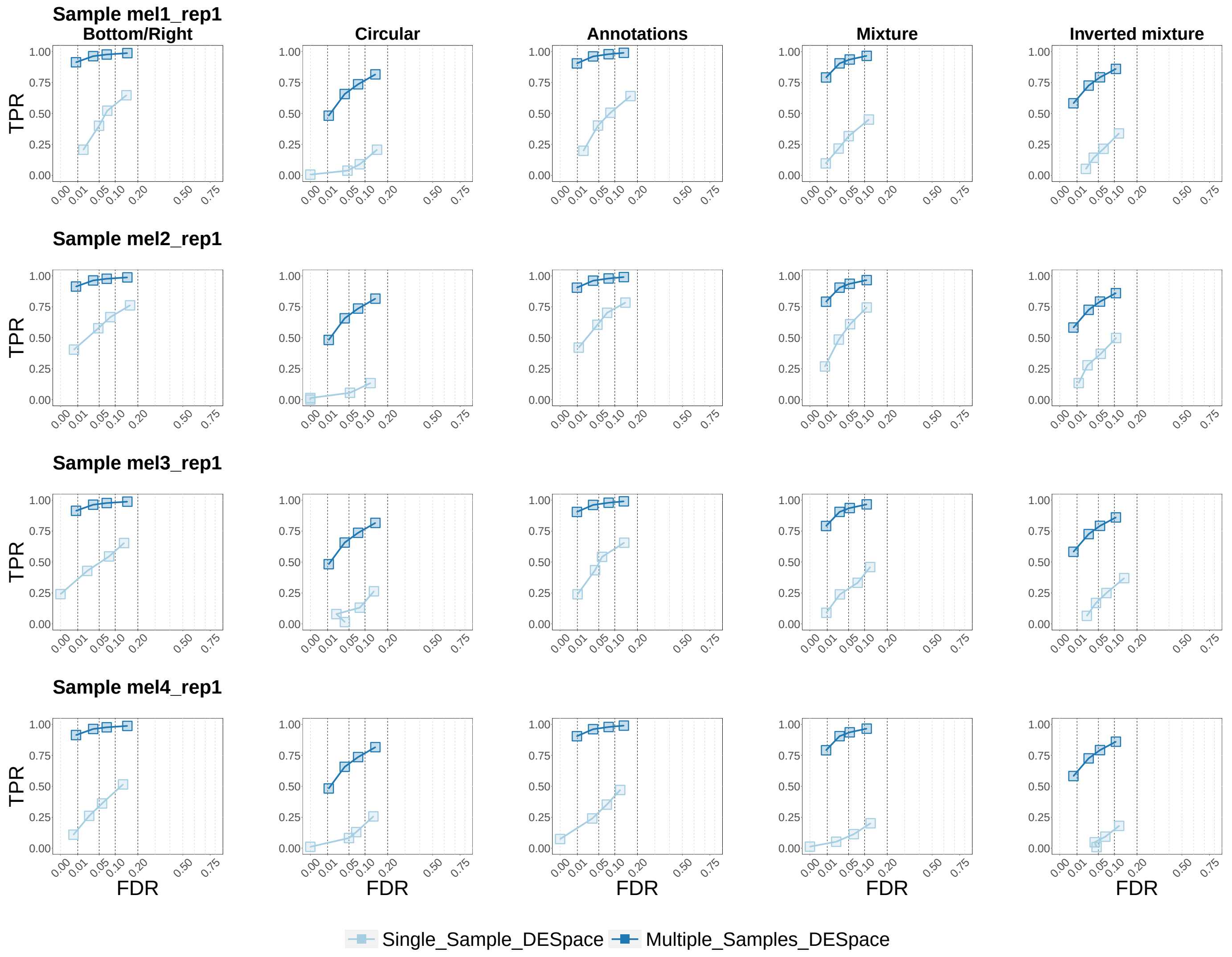


**Supplementary Figure 16:** TPR vs. FDR for SVG detections in the multi-sample simulations, in each *melanoma* sample. Rows and columns refer to the specific sample used as anchor data in the simulation, and to the SV profiles, respectively. *Single_ sample_ DESpace* and *Multiple_ samples_ DESpace* indicate *DESpace* on single-sample and multi-sample modes, respectively.

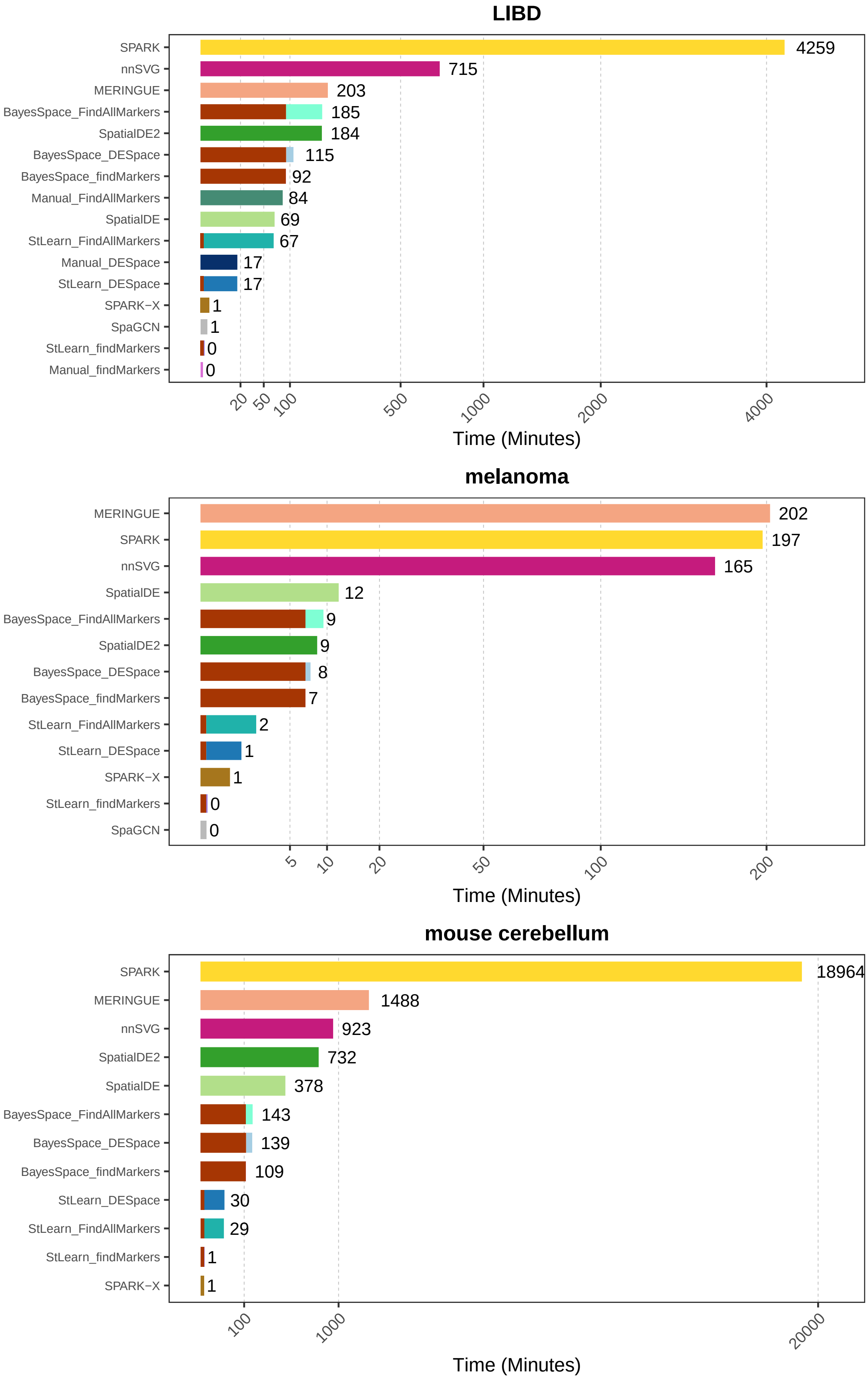


**Supplementary Figure 17:** Runtime (in minutes), of each method, in the real datasets; for the *LIBD* and *melanoma* data, we report the average runtime across replicates. For *DESpace*, *findMarker*, and *FindAllMarkers* we colour differently the runtime to compute spatial clusters (brown left region), and the one to run the differential methods themselves. Note that, for the *LIBD* dataset, *SPARK* successfully ran on Sample 151507 only, hence *SPARK*'s runtime refers to that sample only.

### References


[1] Y. Benjamini and Y. Hochberg. Controlling the false discovery rate: a practical and powerful approach to multiple testing. *Journal of the Royal statistical society: series B (Methodological)*, 57(1):289–300, 1995.

[2] A. T. L. L. Q. F. W. Davis J McCarthy, Kieran R Campbell. Scater: pre-processing, quality control, normalization and visualization of single-cell RNA-seq data in R . *Bioinformatics*, 33(8):1179–1186, 2017.

[3] M. W. Lukas, T. Leonardo, Collado, and C. H. Stephanie. Orchestrating spatially-resolved transcriptomics analysis with bioconductor. https://lmweber.org/OSTA-book/quality-control.html. Accessed: 2022-08-07.

[4] A. T. Lun, D. J. McCarthy, and J. C. Marioni. A step-by-step workflow for low-level analysis of single-cell rna-seq data with bioconductor. *F1000Research*, 5, 2016.

[5] D. Ruben, Z. Qian, D. Rui, C.-H. L. Eng, L. Huipeng, L. Kan, F. Yuntian, Z. Tianxiao, S. Arpan, B. Feng, R. E. George, P. Nico, C. Long, and Y. Guo-Cheng. Giotto: a toolbox for integrative analysis and visualization of spatial expression data. *Genome Biol*, 22:78, 2021.

[6] R. Satija, J. A. Farrell, D. Gennert, A. F. Schier, and A. Regev. Spatial reconstruction of single-cell gene expression data. *Nature biotechnology*, 33(5):495–502, 2015.
